# Long-range inhibitory control of cholinergic network dynamics in the striatum

**DOI:** 10.64898/2026.06.29.735329

**Authors:** Samet Kocaturk, Elif Beyza Guven, James M Tepper, Maxime Assous

## Abstract

Striatal cholinergic interneurons (CINs) exhibit a transient pause in tonic firing in response to salient stimuli, a hallmark of reinforcement learning that becomes synchronized with learning. Although thalamostriatal and dopaminergic inputs have been implicated in this pause, the underlying circuit mechanisms remain unclear. Here, we combine optogenetics, electrophysiology, and genetic approaches to examine inhibitory interactions within the CIN network. Synchronized activation of CINs in striatal slices elicited robust feedback inhibition in CINs, suppressing firing and generating pause-like responses. This inhibition was mediated by GABA_A_ receptors and required β2-containing nicotinic acetylcholine receptors (β2-nAChR), and could be recruited by thalamostriatal activation. Dopamine is not required for this circuit but modulates it via D2 receptors. Surprisingly, cell-type–specific silencing and striatal β2-nAChR deletion excluded local GABAergic sources, whereas retrograde β2-nAChR deletion abolished inhibition, revealing an extrastriatal pathway. These findings identify a long-range inhibitory mechanism linking synchronized cholinergic activity to pause generation in striatal circuits.

## Introduction

The striatum exhibits some of the highest levels of cholinergic markers in the brain, reflecting the prominent role of acetylcholine (ACh) in basal ganglia function^1–3^. This cholinergic tone arises almost entirely from a small population of cholinergic interneurons (CINs), which comprise only ∼1–2% of striatal neurons yet exert widespread influence through their dense axonal arborizations^4,5^. Despite their scarcity, CINs are critical regulators of reward processing, attention, and behavioral flexibility, largely through their ability to modulate synaptic integration and striatal circuit dynamics^6–10^.

A defining feature of CIN activity is their coordinated population response to salient stimuli, often expressed as a transient pause in firing embedded within a multiphasic burst–pause–rebound pattern^11–19^. These pauses are highly synchronized across CINs and are closely linked to reinforcement learning, attentional processes, and behavioural adaptation^15,18,20–22^. However, the cellular and circuit mechanisms underlying both pause generation and synchronization remain incompletely understood.

Several non-mutually exclusive mechanisms have been proposed. Dopamine is widely recognized as a key modulator of CIN activity and pause dynamics. Disruption of dopaminergic signalling alters pause expression, and activation of D2 receptors contributes to shaping their timing, magnitude, and synchronization^15,18,23–27^. However, pauses can persist in the absence of dopamine neuron activity^15,18^, or following dopamine depletion^23^, indicating that dopaminergic signalling alone is not sufficient to account for these dynamics.

Glutamatergic inputs, particularly from the parafascicular nucleus of the thalamus (PfN), provide a powerful excitatory drive onto CINs and are thought to play a central role in synchronizing their activity^14,17,24,28–31^. Thalamostriatal activation can evoke robust multiphasic responses in CINs, with the magnitude of the initial excitation influencing subsequent pause duration^14,17,24^. These observations suggest that convergent excitatory inputs may initiate coordinated activity across CIN populations, although the mechanisms that transform excitation into synchronized inhibition remain unclear.

GABAergic signalling represents another major candidate mechanism. CINs receive a disproportionately high level of inhibitory input^14^ and are innervated by both intrinsic and extrinsic GABAergic sources, including striatal interneurons, projection neurons, and long-range afferents^7,32–36^. In addition, CINs can recruit local GABAergic microcircuits through activation of β2-containing nicotinic acetylcholine receptors (β2-nAChRs), leading to disynaptic inhibition of spiny projection neurons (SPNs,^1,37–41^). Importantly, β2-nAChRs activation can also evoke GABAergic responses onto CINs themselves, supporting the existence of recurrent inhibitory motifs within the cholinergic network^1,33,42,43^.

Consistent with this idea, multi-neuronal recordings have demonstrated that activation of one or a few CINs can evoke polysynaptic inhibition in neighboring CINs via GABAergic intermediaries^1,33,42^. These studies suggest that such coupling may partially involve specific interneuron subtypes, including tyrosine hydroxylase–expressing interneurons (THINs,^33^). Notably, the strength and reliability of these inhibitory interactions increase with the extent of CIN recruitment, indicating that network-level dynamics may differ substantially from unitary connectivity^33,42^.

This distinction is particularly important given that CINs in vivo operate as coordinated populations. Synchronous activation of CINs has profound circuit-level consequences, including the triggering of dopamine and serotonin release via β2-nAChRs on dopaminergic and serotoninergic axons, respectively^44–46^ as well as widespread modulation of striatal output^10,47^. Whether similar population-level mechanisms govern inhibitory interactions among CINs themselves, and how these contribute to pause generation, remains unknown.

Here, we address these questions by examining how synchronized activation of CINs shapes inhibitory signalling within the CIN network. Using optogenetic strategies to recruit either thalamostriatal inputs or CIN populations directly, we show that coordinated CIN activity elicits robust and reliable feedback inhibition capable of interrupting their tonic firing. This inhibition depends on β2-nAChRs and GABA_A_ receptor activation, consistent with recruitment of a GABAergic intermediary. Strikingly, we find that this inhibitory pathway does not require local striatal GABAergic neurons but instead depends on extrastriatal afferents, as demonstrated by selective genetic manipulations. In addition, we show that this circuit is strongly modulated by dopamine through D2 receptor signaling, which suppresses feedback inhibition without being required for its expression. Together, these findings identify a previously unrecognized long-range inhibitory mechanism controlling CIN activity and provide new insight into how synchronized cholinergic signalling generates pause responses and shapes striatal function.

## Results

### Thalamostriatal activation recruits β2-nAChR-dependent feedback inhibition in CINs

Previous studies have characterized inhibitory coupling between CINs using unitary or small-scale activation paradigms^33,42^, providing important insights into local circuit interactions. Here, we examined how synchronized, population-level activation of CINs influences inhibitory coupling. Thalamostriatal input provides a physiologically relevant means to achieve such coordinated activation. PfN afferents constitute a major excitatory drive onto CINs, contribute to the generation of burst–pause responses, and have been shown to effectively synchronize CIN activity both in vivo and in slices^14,24,28^. Under ex vivo conditions, this synchronized activation is sufficient to trigger dopamine release via β2-containing nAChRs expressed on dopaminergic axons^45^. Given that the same class of nAChRs has been implicated in mediating polysynaptic inhibition between CINs, we hypothesized that thalamostriatal activation may similarly recruit and potentiate feedback inhibitory coupling within the CIN network.

To test this, we targeted thalamostriatal projections using injection of a Cre-dependent ChR2 AAV into the PfN of VGLUT2-Cre mice, or injection of a CaMKIIα-ChR2 AAV into the PfN of wild-type mice. We then recorded from CINs in dorsal striatal slices while optically stimulating PfN axons (Fig 1A,B, Fig. S1 A).

**Figure 1.**
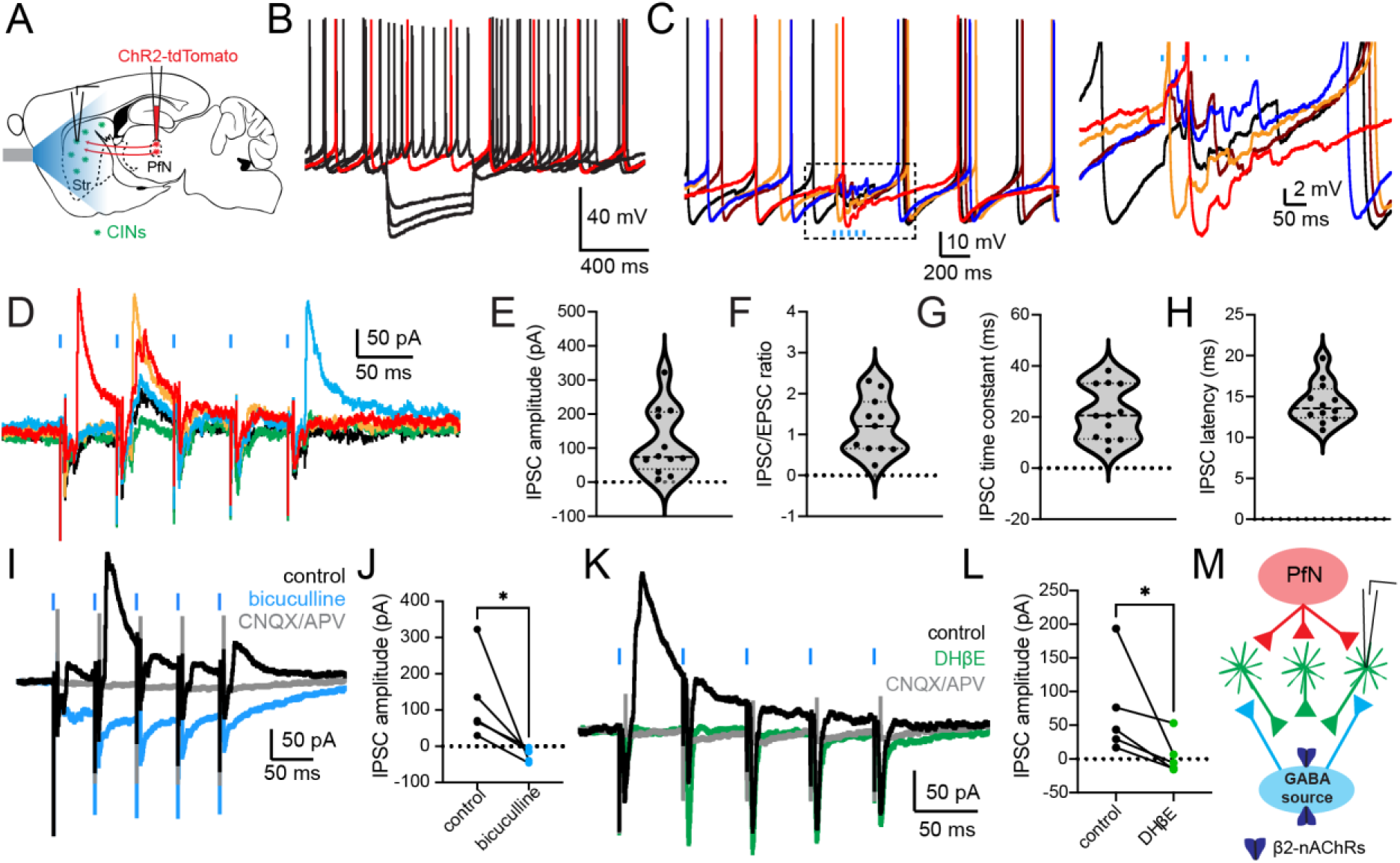
Thalamostriatal input recruits β2-nAChR-dependent GABAergic inhibition in cholinergic interneurons. (A) Schematic of experimental approach: ChR2 expression in PfN neurons of VGLUT2-Cre mice and optical stimulation of thalamostriatal terminals in dorsal striatum. (B) Representative voltage response of a CIN to hyperpolarising and depolarising current steps. (C) Current-clamp recordings from a CIN showing excitation followed by a pause in firing upon PfN stimulation. A zoomed view of the boxed region in the left panel is shown on the right, highlighting the EPSP–IPSP sequence. (D) Voltage-clamp recordings (Vh = –45 mV) showing EPSCs followed by IPSCs in CINs. (E) Quantification of IPSC amplitude evoked by PfN stimulation. (F) Quantification of the IPSC/EPSC ratio measured in CINs exhibiting both excitatory and inhibitory synaptic components. (G) Distribution of IPSC decay time constants. (H) Quantification of IPSC onset latency following optical stimulation of PfN afferents. Each dot represents one recorded cell. Violin plots show data distribution with median and interquartile range indicated by dashed lines. (I) IPSCs are abolished by bicuculline (blue, 10 μM), indicating mediation by GABAA receptors. (J) Quantification of IPSC amplitude under control and bicuculline conditions (p=0.031, Wilcoxon matched-pairs signed rank test, n=5). (K) IPSCs are also abolished by the β2-nAChR antagonist DHβE (green, 1 μM). (L) Quantification of IPSC amplitude under control and DHβE conditions (p=0.03, Wilcoxon matched-pairs signed rank test, n=5). (M) Proposed circuit model: PfN activation excites CINs, which recruit a β2-nAChR-dependent GABAergic source providing feedback inhibition onto CIN.

Consistent with previous studies, brief trains of optical stimulation (5 pulses at 20 Hz) evoked robust excitation in all recorded CINs, frequently inducing action potential firing (Fig. 1 C). In cell-attached and current-clamp recordings, most CINs responded to PfN stimulation with a subsequent pause in firing (Fig. 1 C). In current clamp, inhibitory postsynaptic potentials (IPSPs) were detected in the majority of CINs (17/26 cells), typically following an EPSP or action potential (Fig. 1 C and Fig. S1 B). Although the IPSP generally occurred once per trial, its precise temporal position varied across cells and from trial to trial (Fig. 1C, Fig. S1 C). Similarly, in voltage clamp at Vh = –45 mV, PfN stimulation evoked EPSCs followed by fast IPSCs in most CINs (12/19 cells, Fig. 1 D). In roughly half of responsive cells, the IPSC occurred predominantly after the first optical pulse (6/12), whereas in others it followed the second (4/12), fourth (1/12), or fifth (1/12) pulse (Fig. S1 C).

These PfN-evoked IPSCs were large (115.1 ± 28.1 pA, *n* = 12, Fig. 1 E) and frequently exceeded the concomitant EPSC at this holding potential (IPSC/EPSC ratio: 1.22 ± 0.21, *n* = 11, Fig. 1 F). Their relatively slow decay (τ = 21.6 ± 3.06 ms, *n* = 12, Fig. 1 G) and long latency (14.22 ± 0.72 ms, *n* = 12, Fig. 1 H) were consistent with recruitment of a polysynaptic inhibitory circuit. Accordingly, bath application of the GABA_A_ receptor antagonist bicuculline (10 μM) completely abolished the IPSCs (–148.2 ± 55.4 pA, *p* = 0.031, Wilcoxon matched-pairs signed-rank test, *n* = 5, Fig. 1 I-J). Likewise, the type II nAChR antagonist DHβE (1 μM) also eliminated the PfN-evoked IPSCs (–67.03 ± 30.5 pA, ∼93% reduction, *p* = 0.03, *n* = 5, Fig. 1 K-L). Together, these results indicate that thalamostriatal input can effectively recruit a β2-nAChR-dependent GABAergic pathway that feeds back onto CINs and may contribute to pause generation following salient thalamic drive (Fig. 1 M).

### Population activation of CINs evokes large and reliable polysynaptic feedback currents

To directly examine feedback inhibition recruited by synchronized CIN activation, we recorded from CINs in ChAT-ChR2-eYFP mice, in which CINs natively express channelrhodopsin and can be activated by brief blue light pulses (2 ms, 450 nm, Fig. 2 A-B). Optical stimulation reliably evoked action potentials in recorded CINs. To isolate the recurrent synaptic component from the direct photocurrent, recordings were performed with QX-314 (3 mM) in the internal solution to block voltage-gated Na^+^ channels and suppress action potential generation in the recorded cell. Under these conditions, the remaining photocurrent was followed in the vast majority of CINs by a large postsynaptic current (PSC), often composed of multiple synaptic events (Fig 2 C-D).

**Figure 2.**
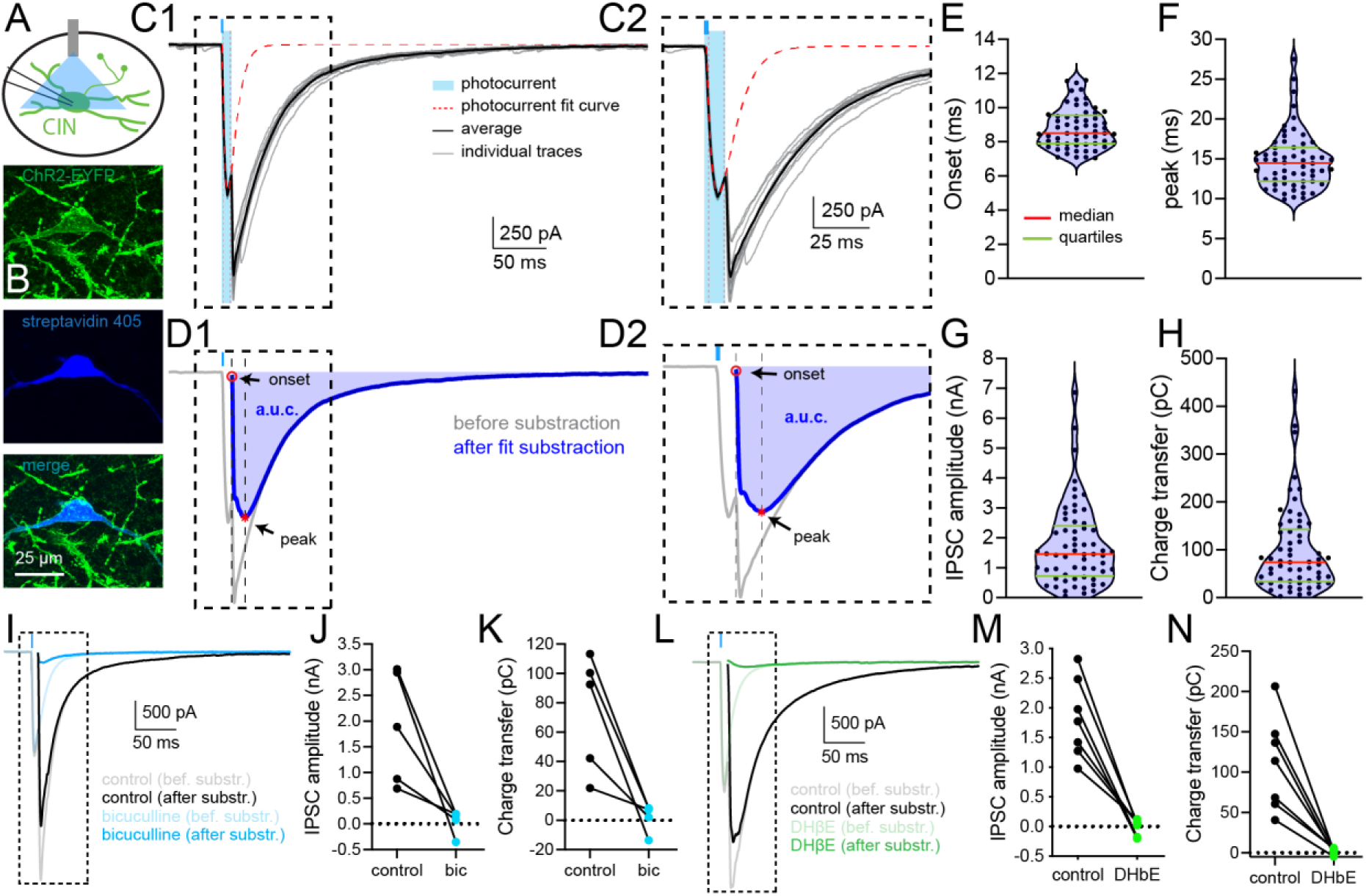
Synchronized CIN activation evokes β2-nAChR-dependent polysynaptic GABAergic currents. (A) Schematic of optogenetic activation of CINs in ChAT-ChR2-eYFP mice. (B) Representative fluorescence images showing ChR2-eYFP expression in biocytin-filled CINs (green), labelled posthoc with streptavidin 405 (blue). (C) Representative voltage-clamp recordings showing light-evoked responses in a CIN (individual trials in grey and average response in black) and the photocurrent fitted curve using a biexponential function (red dotted, validated against TTX data). The area highlighted in C1 is expanded in C2. (D1) This fitted curse was then subtracted from CIN recordings to isolate the synaptic component (blue, after fit substraction). (D2) Expanded views illustrating determination of response onset, peak, and area under the curve (a.u.c.) before (grey) and after (blue) subtraction. (E–H) Quantification of response properties after subtraction, including onset latency (E), peak time (F), IPSC amplitude (G), and charge transfer (H). Violin plots show distribution, median, and quartiles. (I) Representative traces showing light-evoked responses before and after subtraction under control conditions and following bicuculline (10 μM, blue). (J–K) Bicuculline significantly reduces IPSC amplitude and charge transfer, indicating mediation by GABAA receptors. (L) Representative traces before and after subtraction under control conditions and following DHβE (1 μM, green). (M–N) DHβE abolishes IPSC amplitude and charge transfer, demonstrating dependence on β2-containing nicotinic receptors.

Because the photocurrent and recurrent PSC overlapped substantially in time (Fig. 2 C and Fig. S2 A), we used a biexponential EPSC-like function to fit the direct photocurrent and subtract it from the total response (A_i_ x (1 − e^−(t−t0)/τrise^) x e^−(t−t0)/τdecay^, Fig. S2B, see methods). This function was parameterized using constant photocurrent kinetics (τ*_rise_*, τ*_decay_*) with varying amplitude across trials. Fig 2 C, Fig. S2) To validate this approach, the same fitting procedure was applied to recordings obtained in TTX (1 μM), which abolishes synaptic transmission and leaves an isolated photocurrent. Under these conditions, the fit closely matched the photocurrent (R = 0.99), confirming that the subtraction procedure accurately isolates the synaptic component (Fig. S2 B-C).

The isolated PSCs had a relatively long onset latency (8.81 ± 0.15 ms, *n* = 61, Fig 2 E) and peak time (14.81 ± 0.47 ms, *n* = 61, Fig. 2 F), consistent with a polysynaptic origin. They were also large in magnitude, with a mean amplitude of 1742 ± 174.1 pA and charge transfer of 98.32 ± 11.46 pC (*n* = 61, Fig. 2 G-H). Thus, synchronized optogenetic activation of CINs evokes large and highly reliable feedback PSCs in neighboring CINs, consistent with recruitment of a robust inhibitory network.

We next determined the pharmacological identity of the large feedback PSCs recorded directly in ChAT-ChR2-eYFP CINs. Consistent with thalamic stimulation and prior work using single-cell or multi-neuronal recordings^33,42^, these population-evoked feedback PSCs were almost completely abolished by bicuculline (10 μM; amplitude change: –1.82 ± 0.55 nA, ∼97% reduction, *p* = 0.031; charge transfer: –71.68 ± 18.68 pC, ∼96.7% reduction, *p* = 0.03; *n* = 5, Fig. 2 I-K) and by DHβE (1 μM; amplitude change: –1.82 ± 0.26 nA, ∼100% reduction, *p* = 0.015; charge transfer: –108.7 ± 21.94 pC, ∼98.3% reduction, *p* = 0.0025; *n* = 7, Fig. 2 L-N). These results demonstrate that the feedback PSCs are polysynaptic GABAergic currents that require β2-nAChR activation.

Because CINs have been reported to co-release GABA under some conditions, we asked whether a residual GABAergic component might persist after nicotinic blockade. However, when bicuculline was applied after DHβE, it produced no additional reduction in the residual response (Fig. S3 A-B, bicuculline + DHβE vs. DHβE alone; amplitude: +0.005 ± 0.009 nA, *p* = 0.75, Fig. S3 C; charge transfer: +0.13 ± 0.47 pC, *p* > 0.999; *n* = 3, Fig. S3 D), indicating that CIN GABA co-release does not measurably contribute to the polysynaptic feedback IPSCs evoked by CIN population stimulation.

### Feedback inhibition recruited by CIN population activity suppresses firing in neighboring CINs

To determine how this inhibitory circuitry influences CIN firing activity while avoiding direct ChR2-mediated photocurrents in the recorded cell, we used a Cre/Flp intersectional viral strategy in ChAT-Cre mice (*N* = 17 mice). In this approach, AAV5-Con/Foff-ChR2-eYFP drove ChR2-eYFP expression selectively in Cre+/Flp- CINs, whereas AAV5-DIO-Flp and AAV5-Flp-mCherry induced Flp and mCherry expression in a subset of CINs, thereby switching off ChR2 in those neurons. This generated two distinct CIN populations within the same striatum: ChR2-eYFP+ CINs that could be optogenetically activated and mCherry+/ChR2-eYFP- CINs that could be recorded without direct photocurrent contamination (Fig. 3 A-C, Fig S4 A).

**Figure 3.**
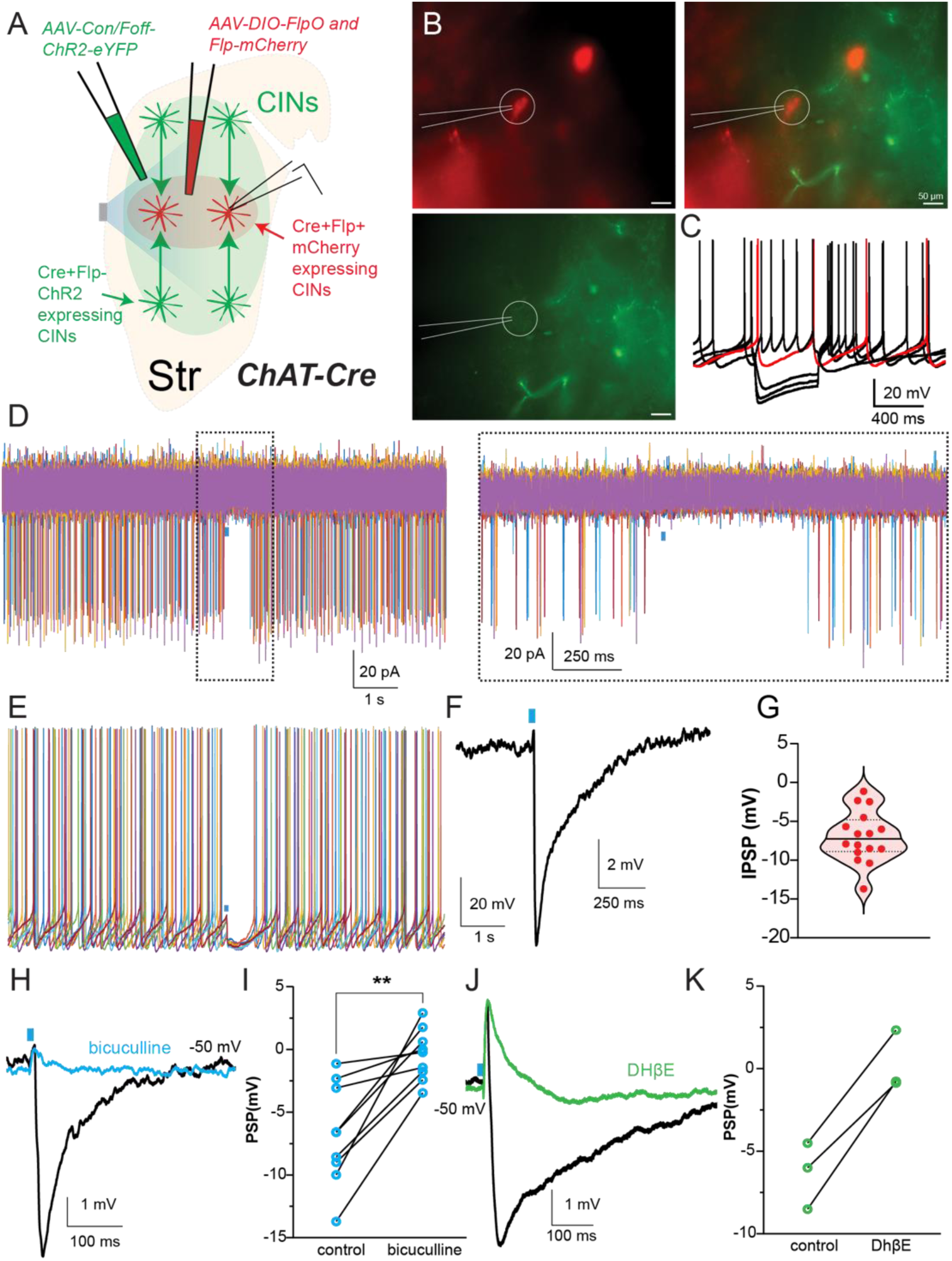
Synchronized activation of CINs suppresses firing in neighboring CINs. (A) Intersectional viral strategy in ChAT-Cre mice to generate distinct populations of ChR2-expressing (Con-Foff-ChR2+) and non-expressing (ChR2−, mCherry+) CINs. (B) Representative fluorescence images showing a recorded mCherry-expressing, ChR2- CIN in striatal slices. (C) Characteristic voltage response of a CIN to negative and positive somatic current injection. (D) Cell-attached recordings showing pause-like suppression of firing in ChR2−/mCherry+ CINs following activation of neighboring CINs (ChR2+, 450nm, 2ms pulse). The boxed region is expanded on the right panel. (E) Current-clamp recordings from ChR2−/mCherry+ CINs showing pause-like suppression of spontaneous firing following activation of neighboring CINs. (F) Current-clamp recordings from hyperpolarized CINs showing inhibitory postsynaptic potentials (IPSPs) evoked by CIN population activation. (G) Quantification of IPSP amplitude measured in CINs. (H) IPSPs recorded in ChR2−/mCherry+ CINs under control conditions and following bath application of bicuculline (10 μM). (I) Quantification of IPSP amplitude reduction following bicuculline. (J) IPSPs recorded in ChR2- /mCherry+ CINs under control conditions and following DHβE (1 μM). (K) Quantification of IPSP amplitude reduction following DHβE.

Using this strategy, we found that synchronized activation of ChR2-eYFP^+^CINs induced clear pause-like responses in neighboring mCherry^+^/ChR2^-^ CINs (Fig. 3 D). In whole-cell current clamp, optical stimulation suppressed spontaneous spiking in 7/8 recorded CINs. Brief (2 ms) optical stimulation also evoked large synaptic inhibitory responses in these neighboring CINs, including IPSCs in voltage clamp (149.8 ± 21.28 pA, *n* = 13, Fig. S4 B-C) and IPSPs in current clamp (–6.96 ± 0.82 mV, *n* = 16, Fig. 3 E-G). These findings demonstrate that synchronized activation of CINs is sufficient to engage inhibitory signaling that suppresses spiking in neighboring CINs, consistent with a polysynaptic inhibitory interaction within the cholinergic network.

We next confirmed the synaptic nature of the inhibition recorded in neighboring mCherry^+^/ChR2^-^ CINs (Fig. 3 H-K, Fig. S4 B-C). These IPSCs were abolished by the GABA_A_ receptor antagonist bicuculline (control: 142.5 ± 34.62 pA vs.10 μM bicuculline; –21.09 ± 9.61 pA, *n* = 7, *p* = 0.016, Wilcoxon matched-pairs test; Fig. S4 B,C). Similarly, the β2-nAChR antagonist DHβE (1 μM) eliminated light-evoked IPSCs (control: 160.5 ± 32.74 pA; DHβE: – 1.67 ± 3.66 pA, *n* = 4; Fig. S4 D,E).

Similarly, in current clamp, IPSPs (–6.77 ± 1.35 mV) were markedly reduced by bicuculline (–0.44 ± 0.68 mV, *n* = 9, *p* = 0.004; Fig. 3 H,I) and DHβE (control: –6.34 ± 1.17 mV; DHβE: 0.24 ± 1.05 mV, *n* = 3; Fig. 3 J,K). Together, these data indicate that synchronized activation of CINs recruits a β2-nAChR-expressing GABAergic source that mediates GABA_A_ receptor-dependent inhibition of neighboring CINs.

### Dopamine depletion does not abolish feedback inhibition among CINs

Because dopamine is known to regulate CIN excitability, notably via D2 receptors, we next asked whether dopaminergic signaling contributes to the feedback inhibition among CINs. We used unilateral 6-OHDA lesions in ChAT-ChR2-eYFP mice to induce extensive loss of nigrostriatal dopaminergic terminals (Fig. 4 A-C). If dopamine release were required to sustain this inhibitory circuit, dopamine depletion should reduce CIN-evoked feedback IPSCs.

**Figure 4.**
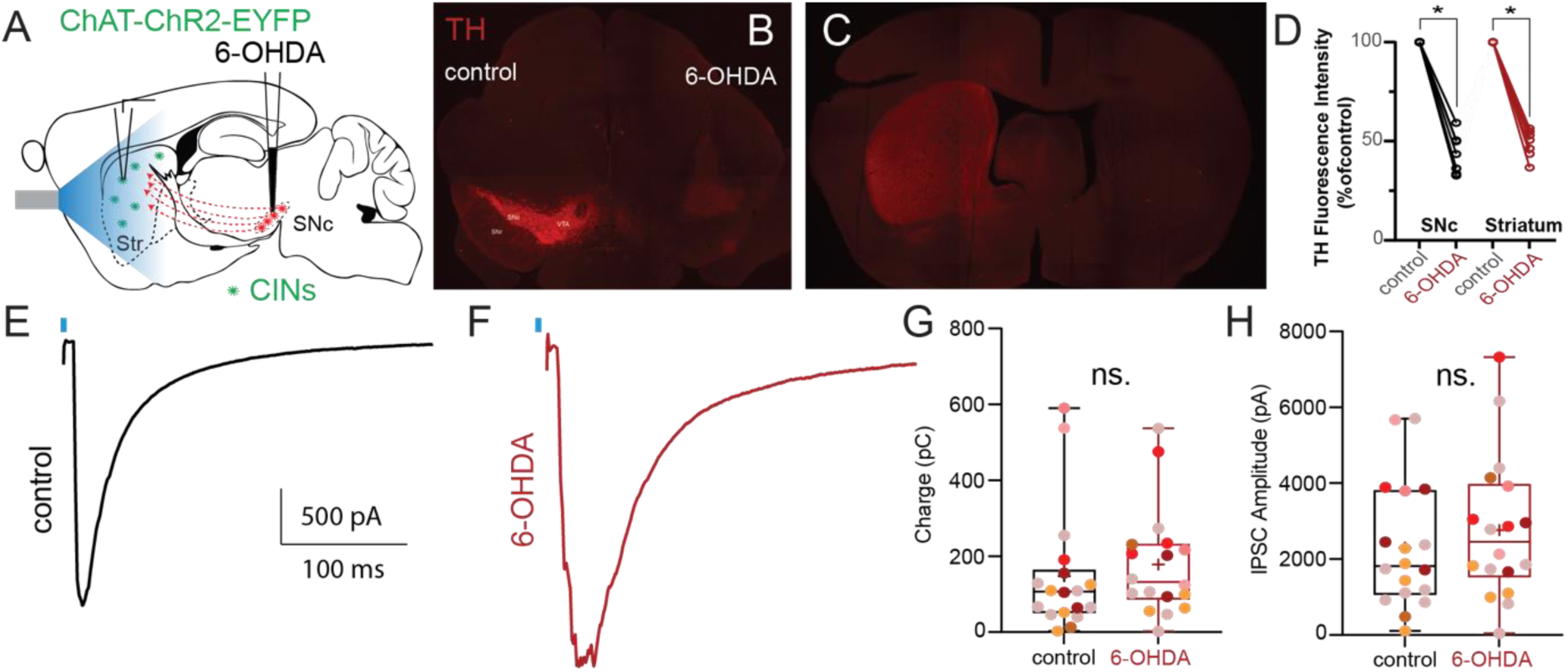
Feedback inhibition of CINs is preserved following dopamine depletion. (A) Schematic of 6-OHDA lesion of dopaminergic neurons in ChAT-ChR2-eYFP mice. (B–C) Representative photomicrographs of tyrosine hydroxylase (TH) immunostaining following unilateral 6-OHDA injection in the SNc. These show unilateral loss of TH immunostaining in SNc (B) and striatum (C). (D) Quantification of TH fluorescence intensity in SNc and striatum, normalized to control. (E) Representative IPSCs recorded from CINs in control striatum following optogenetic activation of CINs (2ms, 450nm). (F) Representative IPSCs recorded from CINs with the same stimulation protocol in 6-OHDA-lesioned striatum. (G–H) Quantification of IPSC charge transfer (G) and amplitude (H) in control and 6-OHDA conditions. Each point represents a cell; each color represent a different mouse. No significant differences were observed (amplitude: p = 0.4064; charge transfer: p = 0.3231, Mann–Whitney U test).

However, this was not the case. In control striatum, population activation of CINs induced robust feedback IPSCs (2304.0 ± 389.6 pA, charge: 147.8 ± 38.66 pC, *N* = 7 mice, *n* = 18; Fig. 4 E,G,H). In the 6-OHDA-treated hemisphere, IPSCs were not significantly altered (2766.0 ± 442.1 pA, *p* = 0.4064; charge: 178.5 ± 33.34 pC, *p* = 0.3231; Mann–Whitney U test; *n* = 18; Fig. 4 F-H). Thus, chronic depletion of nigrostriatal dopamine does not abolish feedback inhibition among CINs, indicating that this circuit is largely preserved after dopamine loss.

### Dopamine receptor signaling acutely modulates feedback inhibition, with a major role for D2 receptors

We next examined whether dopamine receptor manipulations modulate CIN-evoked IPSCs under baseline conditions.

Bath application of the combined D1 and D2 receptor antagonists SCH-23390 (10 μM) and sulpiride (5 μM) significantly reduced both IPSC amplitude (control: 2194.0 ± 443.9 pA; SCH-23390 + sulpiride: 1410.0 ± 297.7 pA; *n* = 7; *p* = 0.0156) and charge transfer (control: 109.5 ± 21.18 pC; antagonist: 68.3 ± 13.27 pC; *p* = 0.0156; Fig. 5 A,B). Likewise, co-application of the D1 and D2 receptor agonists SKF-38393 (10 μM) and quinpirole (10 μM) also attenuated the feedback IPSC (control: 4068.0 ± 798.4 pA vs. agonists: 651.8 ± 533.2 pA; control charge: 194.2 ± 52.15 pC vs. agonists: 35.82 ± 29.76 pC; *n* = 6; *p* = 0.0312; Fig. S5 A,B). These results suggest that feedback inhibition among CINs is maintained within a relatively narrow dopaminergic operating range, such that both reduced and excessive receptor activation weaken the inhibitory response.

**Figure 5.**
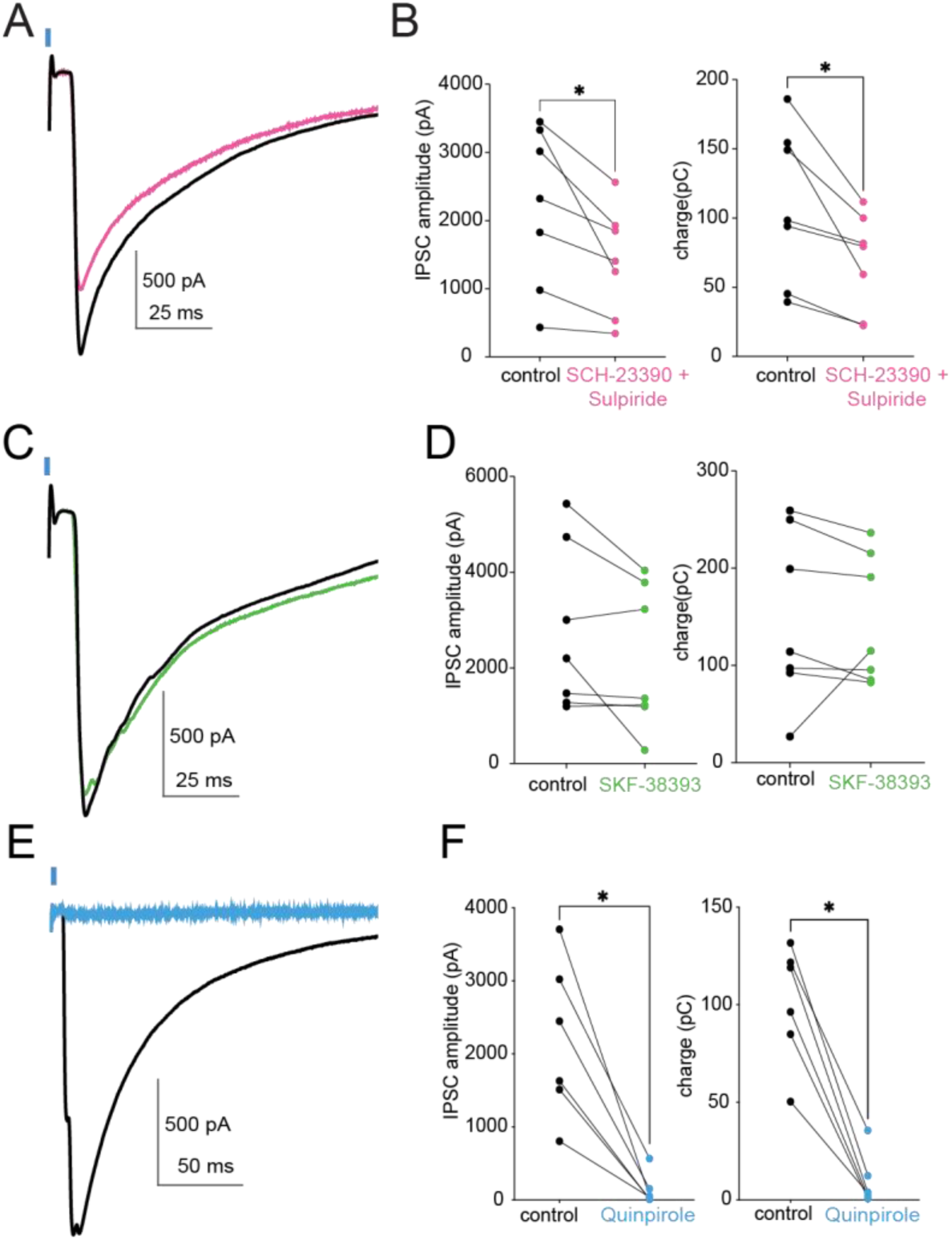
Dopamine bidirectionally modulates feedback inhibition between cholinergic interneurons through D2 receptor signaling. (A) Representative voltage-clamp recordings from ChR2-expressing CINs showing optogenetically-evoked feedback IPSCs under control conditions and following co-application of the D1 receptor antagonist SCH-23390 (10 μM) and the D2 receptor antagonist sulpiride (5 μM). (B) Quantification of IPSC peak amplitude and charge transfer before and after dopamine receptor blockade. Combined D1/D2 receptor antagonism significantly reduced feedback inhibitory responses. (C) Representative recordings showing the effect of the D1 receptor agonist SKF-38393 (10 μM) on optogenetically-evoked feedback IPSCs. (D) Quantification of IPSC peak amplitude and charge transfer following SKF-38393 application. D1 receptor activation did not significantly alter feedback inhibition. (E) Representative recordings showing the effect of the D2 receptor agonist quinpirole (10 μM) on feedback IPSCs. Quinpirole nearly abolished the inhibitory response. (F) Quantification of IPSC peak amplitude and charge transfer before and after quinpirole application. D2 receptor activation strongly suppressed feedback inhibition between CINs. Each line represents one recorded cell. *p < 0.05.

To determine whether one receptor class predominates, we next applied the agonists separately. The selective D1 receptor agonist SKF-38393 did not significantly alter IPSC amplitude (control: 2759.0 ± 649.9 pA; SKF-38393: 2160.0 ± 562.4 pA; *p* = 0.1562, Fig. 5C-D) or charge transfer (control: 145.8 ± 24.99 pC; SKF-38393: 148.4 ± 33.39 pC; *p* = 0.297; *n* = 7; Fig. 5 C,D). In contrast, the D2 receptor agonist quinpirole strongly suppressed the inhibitory response (control: 2188.0 ± 437.7 pA; quinpirole: 141.5 ± 87.82 pA; control charge: 100.7 ± 12.29 pC; quinpirole: 9.78 ± 5.45 pC; *n* = 6; *p* = 0.0312; Fig. 5 E,F). Thus, D2 receptor activation, but not D1 receptor activation, potently suppresses feedback inhibition among CINs, indicating that dopaminergic regulation of this circuit is predominantly mediated by D2-dependent signaling.

### Striatal GABAergic neurons are not required for feedback inhibition among CINs

To determine the source of the GABAergic input mediating the feedback IPSCs, we combined optogenetic excitation of CINs with optogenetic silencing of candidate striatal GABAergic populations using halorhodopsin (HaloR3.0, Fig. 6, Fig. S6). On alternating trials, CINs were activated with a blue light pulse (2 ms), either alone or together with a yellow light pulse (700 ms, beginning 200 ms before blue light onset) to silence the targeted population. As a control, yellow light alone had no effect on feedback IPSCs in ChAT-ChR2-eYFP mice (amplitude: – 3.58 ± 36.6 pA, *p* = 0.92; charge transfer: +0.47 ± 1.76 pC, *p* = 0.79; *n* = 12, not shown).

**Figure 6.**
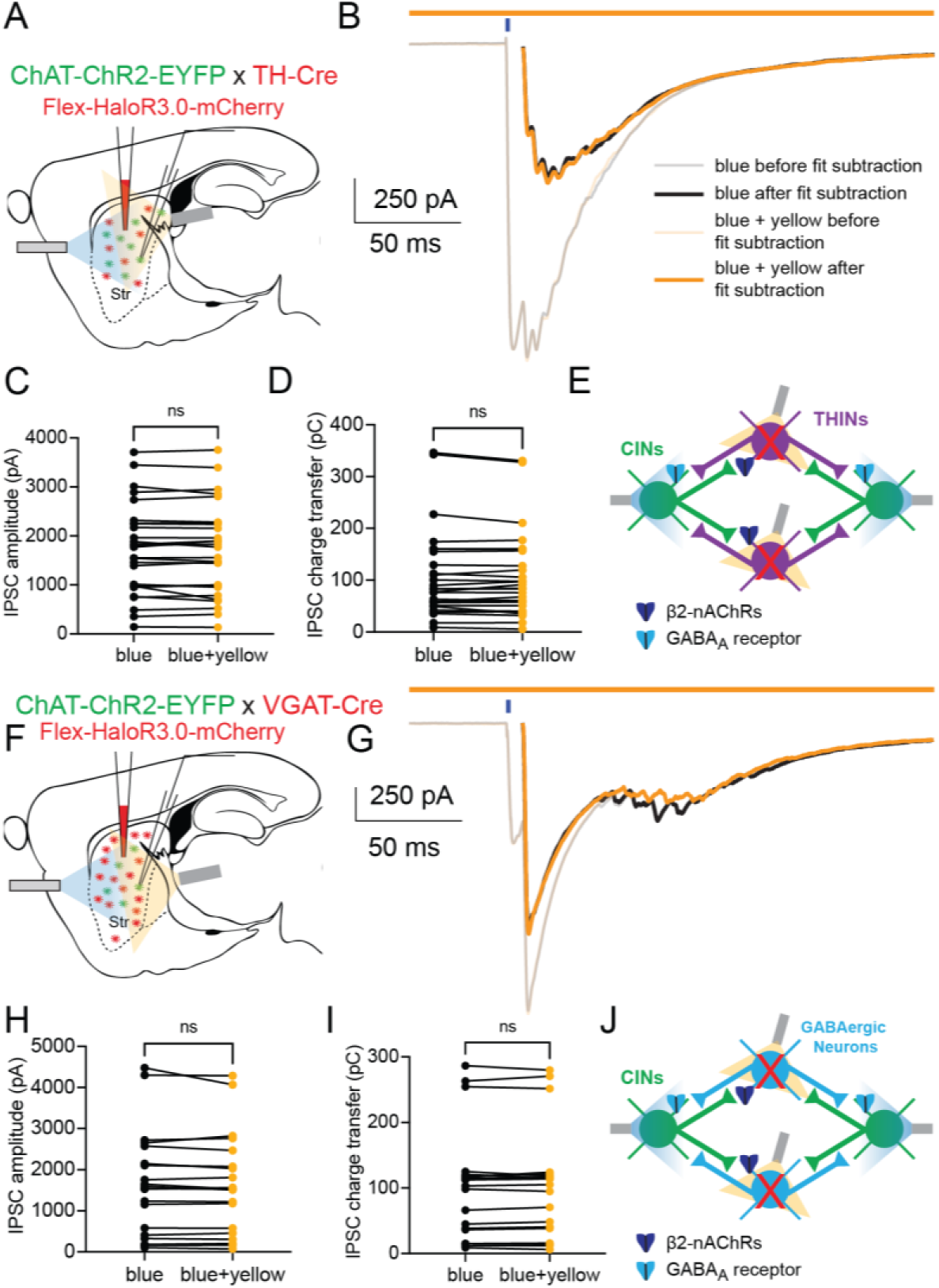
Local striatal GABAergic neurons are not required for feedback inhibition between cholinergic interneurons. (A) Schematic of the double optogenetic strategy used to test the contribution of THINs to feedback inhibition between CINs. ChAT-ChR2-eYFP × TH-Cre mice received striatal injections of a Cre-dependent HaloR3.0-mCherry AAV, enabling optogenetic inhibition of THINs during blue light activation of ChR2-expressing CINs. (B) Representative voltage-clamp recordings from CINs showing optogenetically-evoked feedback IPSCs during blue light stimulation alone and during simultaneous blue + yellow light stimulation to inhibit THINs. Traces are shown before and after photocurrent subtraction. (C,D) Quantification of IPSC peak amplitude (C) and charge transfer (D) under blue light stimulation alone or during simultaneous inhibition of THINs. Optogenetic suppression of THINs did not significantly alter feedback inhibition between CINs. (E) Schematic summary illustrating that THIN inhibition does not disrupt β2-nAChR-dependent polysynaptic feedback inhibition onto CINs. (F) Schematic of the experimental strategy used to broadly inhibit striatal GABAergic neurons. ChAT-ChR2-eYFP × VGAT-Cre mice received striatal injections of a Cre-dependent HaloR3.0-mCherry AAV, allowing optogenetic silencing of GABAergic neurons during activation of CINs. (G) Representative voltage-clamp recordings showing optogenetically-evoked feedback IPSCs during blue light stimulation alone and during simultaneous blue + yellow light stimulation to inhibit striatal GABAergic neurons. Traces are shown before and after photocurrent subtraction. (H,I) Quantification of IPSC peak amplitude (H) and charge transfer (I) under both conditions. Broad inhibition of striatal GABAergic neurons did not significantly affect feedback inhibition between CINs. (J) Schematic summary illustrating that suppression of local striatal GABAergic neurons fails to disrupt β2-nAChR-dependent inhibitory coupling between CINs. Each line represents one recorded cell. ns, not significant.

We first examined THINs, as a recent study implicated THINs in recurrent inhibition following unitary CIN activation. In ChAT-ChR2-eYFP;TH-Cre mice expressing HaloR in THINs (Fig. 6 A), silencing THINs did not alter the feedback IPSC evoked in CINs (control amplitude: 1.72 ± 0.18 nA; change with THIN silencing: –8.3 ± 15.9 pA, *p* = 0.6; charge change: –1.37 ± 1.44 pC, *p* = 0.35; *n* = 27, Fig. 6 B-E). We next tested somatostatin-expressing low threshold spike interneurons (LTSIs) in ChAT-ChR2-eYFP;SST-Cre mice (Fig. S6 A) and found no significant effect of LTSI inhibition (amplitude change: –20.61 ± 33.7 pA, *p* = 0.55; charge change: –2.87 ± 1.9 pC, *p* = 0.16; *n* = 13, Fig. S6 B-D).

To target additional interneuron subclasses, we used ChAT-ChR2-eYFP;Htr3a-Cre mice, in which HaloR expression captures fast spiking interneurons (FSIs), neurogliaform interneurons (NGFs), fast adapting interneurons (FAIs), and spontaneously active bursty interneurons (^38,48,49^, SABIs, Fig. S6 E). Again, optogenetic silencing of Htr3a+ interneurons failed to alter the feedback IPSC (amplitude change: +5.92 ± 40.8 pA, *p* = 0.89; charge change: –1.42 ± 1.68 pC, *p* = 0.41; *n* = 19, Fig. S6 F-H). Finally, in ChAT-ChR2-eYFP;VGAT-Cre mice, broad silencing of striatal GABAergic neurons (Fig. 6 F) also failed to affect feedback IPSCs (amplitude change: –39.8 ± 45.4 pA, *p* = 0.4; charge change: +0.33 ± 1.6 pC, *p* = 0.84; *n* = 11, Fig. 6 G-J). Together, these experiments indicate that the major known classes of striatal GABAergic neurons are not required for the population-induced feedback inhibition of CINs (Fig. 6 J).

To test this conclusion genetically, we conditionally deleted β2-nAChRs from striatal neurons in ChAT-ChR2-eYFP;β2^fl/fl^ mice by unilaterally injecting AAV-hSyn-mCherry-Cre into the striatum, with the contralateral hemisphere receiving AAV-hSyn-mCherry as a control (Fig. 7 A-C). Consistent with the optogenetic disconnection experiments, deleting β2-nAChRs from striatal neurons did not reduce the feedback IPSC (amplitude change: +122.5 ± 474.1 pA, *p* = 0.8; charge change: –2.2 ± 25.2 pC, *p* = 0.93; *n* = 25 Cre, *n* = 17 control, Fig. 7 D-G). Altogether, these findings strongly argue against a major role for intrastriatal GABAergic neurons in this circuit and instead suggest an extrastriatal source.

**Figure 7.**
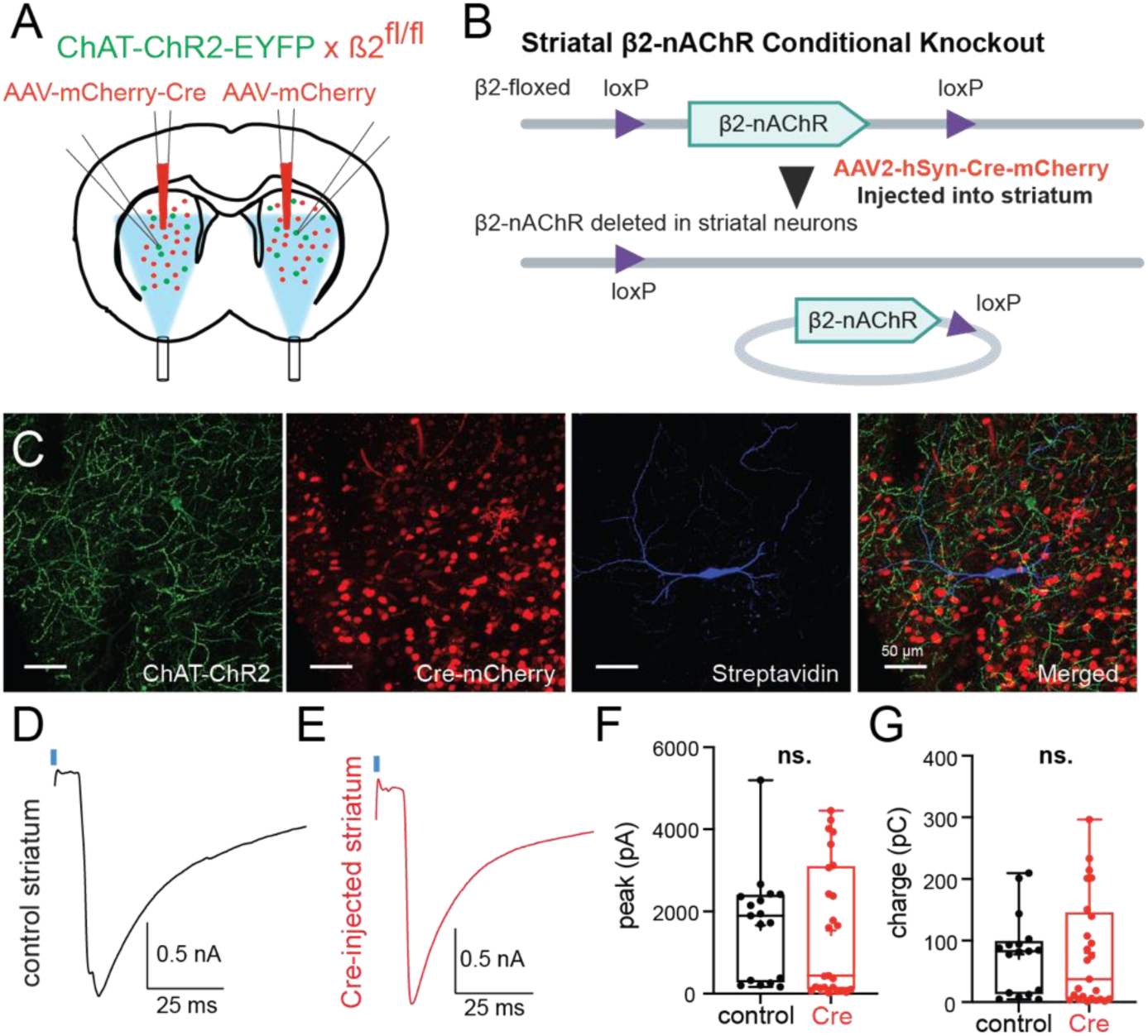
Feedback inhibition persists after deletion of β2-nAChRs from striatal neurons. (A) Experimental design for unilateral Cre-mediated deletion of β2-nAChRs in the striatum of ChAT-ChR2-EYFP × β2fl/fl mice. (B) Schematic of conditional knockout strategy: Cre expression in striatal neurons (via AAV delivery) excises the floxed β2-nAChR gene. (C) Representative histological images showing ChR2-eYFP expression in CINs and Cre-mCherry expression in the injected striatum. CIN was filled with biocytin, revealed with streptavidin 405. (D–E) Representative voltage-clamp recordings showing light-evoked feedback IPSCs in control (D) and Cre-injected (E) striatum. (F–G) Quantification of IPSC peak amplitude (F) and charge transfer (G) showing no significant difference between conditions.

### Feedback inhibition among CINs depends on β2-nAChRs expressed by striatal afferents

To directly test whether the inhibitory feedback among CINs originates from an extrastriatal GABAergic population expressing β2-nAChRs, we used ChAT-ChR2 × β2^fl/fl^ mice and deleted β2-nAChRs in striatal afferents (Fig. 8 A). A viral cocktail containing AAV5-retro-Cre and AAV5-Flex-tdTomato was injected unilaterally into dorsal striatum, while the contralateral hemisphere received a control cocktail of AAV5-retro-Flp and AAV5-Flp-mCherry (*N* = 6 mice, Fig. 8 A-B). This strategy produced Cre-mediated excision of the floxed β2-nAChR gene in neurons projecting to the striatum, thereby removing β2-nAChRs from striatal afferents.

**Figure 8.**
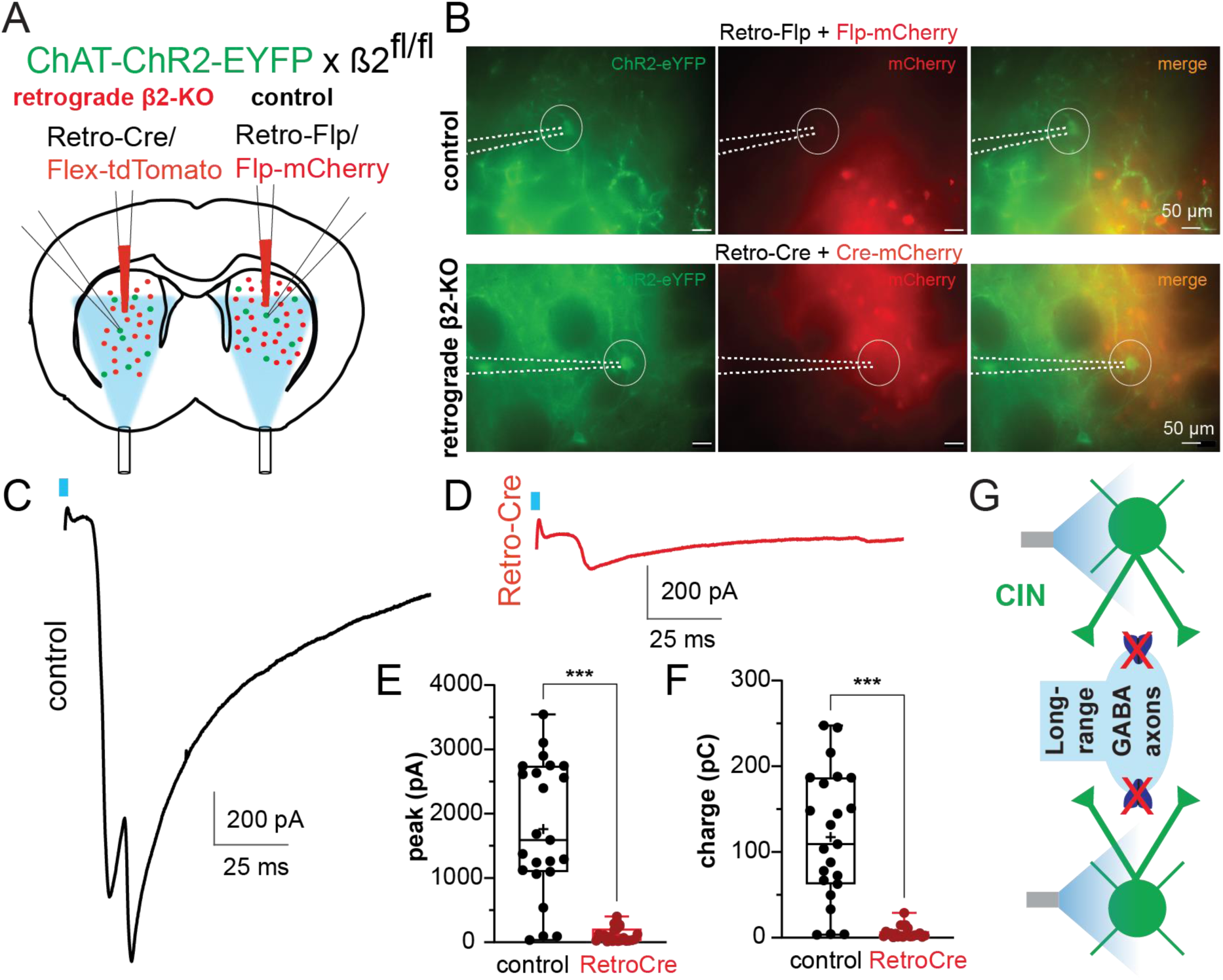
Feedback inhibition among CINs requires β2-nAChRs on extrastriatal afferents. (A) Experimental design for retrograde deletion of β2-nAChRs from striatal afferents in ChAT-ChR2-EYFP × β2fl/fl mice. One hemisphere received Retro-Flp + Flp-mCherry (control), and the other received Retro-Cre + Flex-tdTomato to delete β2-nAChRs in neurons projecting to the striatum. (B) Representative striatal fluorescence images showing recorded ChR2-EYFP-expressing CINs surrounded by mCherry labelled neurons in control and retrograde β2-KO conditions. (C–D) Representative voltage-clamp recordings showing robust light-evoked feedback IPSCs in control striatum (C, retro-Flp) and markedly reduced responses after retrograde β2 deletion (D, retro-Cre). (E–F) Quantification of IPSC peak amplitude (E) and charge transfer (F) showing a significant reduction in the retrograde β2-KO condition (retro-Cre, red). (G) Summary model showing that β2-nAChRs on long-range GABAergic afferents are required for feedback inhibition among CINs.

Whole-cell voltage-clamp recordings from ChR2^+^ CINs showed that brief optical activation of CINs (2 ms, 450 nm) reliably evoked large feedback IPSCs in the control hemisphere (1760.0 ± 218.0 pA, charge: 117.4 ± 15.73 pC, *n* = 23; Fig. 8 C). In striking contrast, deleting β2-nAChRs from striatal afferents markedly suppressed these responses in the retro-Cre hemisphere (122.1 ± 24.47 pA, *p* < 0.001; charge: 5.79 ± 1.42 pC, *p* < 0.001; Mann–Whitney U test; *n* = 23; Fig. 8D-F). These data indicate that β2-nAChRs expressed by striatal afferents are essential for mediating the polysynaptic inhibitory feedback among CINs. Thus, the feedback inhibition observed after CIN population activation arises from an extrastriatal GABAergic population that provides GABA_A_ receptor-mediated inhibition onto CINs (Fig. 8 G).

## Discussion

CINs exhibit a characteristic pause in firing in response to salient stimuli, a signal closely linked to reinforcement learning, attention, and behavioral flexibility^13,19,22,50^. Although this pause has been extensively described in vivo, the circuit mechanisms underlying its generation and synchronization across CIN populations remain incompletely understood. Here, we identify a robust mechanism of polysynaptic inhibition among CINs that emerges during coordinated population activity and depends critically on β2-nAChRs and extrastriatal GABAergic inputs.

### Population-level activation reveals a robust inhibitory architecture among CINs

By mimicking synchronized activation of CINs, we show that inhibitory coupling among CINs becomes a robust network-level phenomenon. Earlier studies using multi-neuron recordings established the existence of CIN-to-CIN polysynaptic inhibition and characterized key unitary properties of this connectivity^33,42^. Our data build directly on that work by demonstrating that when CINs are co-activated, as occurs during behaviorally relevant population events, this coupling scales into a strong and reliable inhibitory mechanism.

Such scaling is highly relevant to in vivo physiology, where CINs operate as coordinated ensembles rather than independent units^20–22^. Under these conditions, feedback inhibition was sufficient to suppress spontaneous CIN firing, suggesting that CINs engage a self-regulating inhibitory loop.

The recruitment of similar inhibitory responses by thalamostriatal stimulation further supports the physiological relevance of this mechanism. The PfN of the thalamus provides a major excitatory drive to CINs and plays a central role in shaping their burst–pause responses^14,24,28,30^. Our results suggest that thalamic excitation not only drives CIN firing but also engages a delayed inhibitory pathway that transforms excitation into coordinated suppression. In this framework, the CIN pause is not simply the absence of excitation or the result of intrinsic adaptation, although intrinsic conductances likely contribute to pause dynamics^19,22,51–54^, but may also reflect an active network process in which synchronized cholinergic activation recruits inhibition onto the CIN population.

### β2-containing nAChRs gate a polysynaptic inhibitory pathway

The dependence of this inhibition on β2-nAChRs places it within a broader framework in which striatal nAChRs act as strategically positioned regulators of transmitter release. While muscarinic receptors are widely expressed and mediate slower modulatory effects^55–57^, nicotinic receptors are more selectively localized, particularly at presynaptic terminals and interneuron populations, enabling rapid and temporally precise control of circuit activity^1,58–60^. This principle is well established for monoaminergic signaling. CIN activation can trigger dopamine release through β2-containing nAChRs on dopaminergic axons independently of midbrain firing^44,45,61,62^. More recently, similar mechanisms have been shown to regulate serotonin release in the striatum^46^. Together, these findings highlight a general role for nAChRs as fast presynaptic control points for transmitter release.

Our results extend this framework to inhibitory signaling. Pharmacological blockade of β2-containing nAChRs or GABA_A_ receptors abolished the feedback response, indicating that synchronized CIN activation recruits a β2-nAChR-dependent GABAergic pathway. This is consistent with prior work showing that CINs engage specific GABAergic interneurons populations to mediate disynaptic inhibition of spiny projection neurons^37–39^. Importantly, the same signalling logic where CINs drive GABA release through the activation of β2-nAChRs is here engaged to inhibit CINs themselves^33,42^.

The timing and pharmacology of the response further support a polysynaptic mechanism and argue against alternative explanations such as direct GABA co-release from CINs, which has been reported in other contexts^63^. Moreover, bicuculline did not further reduce responses after DHβE, arguing against a major contribution of direct GABA co-release from CINs under these conditions. Thus, the most likely interpretation is that CIN-derived acetylcholine activates β2-containing nAChRs on a GABAergic element, which in turn releases GABA onto CINs.

### Local striatal GABAergic circuits are not required

Previous studies have suggested that feedback inhibition among CINs may involve recruitment of local striatal GABAergic interneurons. Early work demonstrated that activation of individual CINs can evoke polysynaptic inhibition in neighboring CINs, with pharmacological evidence implicating nicotinic receptor–dependent engagement of GABAergic intermediaries^42^. Building on this, Dorst et al. (2020) systematically tested the contribution of multiple interneuron populations, including SST-, PV-, Htr3a-, and TH-expressing interneurons, using a combination of optogenetic and chemogenetic approaches. These experiments revealed a partial contribution of THINs.

In this context, a major and unexpected finding of the present study is the lack of involvement of local striatal GABAergic circuits in feedback inhibition among CINs during population activation. This conclusion is supported by converging lines of evidence. First, optogenetic silencing of THINs failed to reduce feedback inhibition, despite their previously reported contribution and known sensitivity to cholinergic signaling^39^. Second, inhibition of broader interneuron populations, including Htr3a-expressing interneurons and virtually all striatal GABAergic neurons using VGAT-Cre mice, similarly had no effect. Notably, these same interneuron populations have been shown to play a central role in mediating disynaptic inhibition of SPNs following CIN activation^38,39^, highlighting a clear dissociation between the circuits controlling SPNs and those mediating CIN feedback inhibition.

Crucially, these optogenetic findings were reinforced by a complementary genetic strategy. Conditional deletion of β2-containing nAChRs from striatal neurons, achieved via local viral delivery of Cre recombinase in β2^fl/fl^ mice^64^, did not alter the feedback inhibitory responses. Given that β2-containing nAChRs are required for CIN-driven recruitment of striatal interneurons, this result provides strong causal evidence that local striatal neurons are not the primary source of the GABAergic inhibition observed here.

One likely explanation for the discrepancy with previous studies is the difference in the scale and pattern of CIN activation. While earlier work probed unitary interactions between small numbers of CINs^33^, the present study examined the consequences of synchronized population activity. Under these circumstances, the circuit appears to preferentially recruit a distinct, non-local inhibitory pathway that dominates over local microcircuit contributions.

More broadly, these findings suggest that the organization of inhibitory control within the striatum is state-dependent and hierarchical, with local interneurons mediating well-established disynaptic inhibition of SPNs, while CIN feedback inhibition engages a separate, long-range GABAergic mechanism. This distinction refines current models of striatal circuitry by demonstrating that nicotinic recruitment of inhibition is not confined to local microcircuits, but can instead access distributed inhibitory networks to regulate cholinergic activity itself. This distinction may be especially important for interpreting CIN function in vivo, where behaviorally relevant events often involve coordinated activity across many CINs rather than isolated unitary interactions^13,15,18,22^.

### Extrastriatal GABAergic inputs mediate feedback inhibition

The evidence for an extrastriatal source of inhibition comes from the retrograde deletion of β2-containing nAChRs in neurons projecting to the striatum, which abolished feedback inhibition. This result demonstrates that the β2-nAChRs critical for the polysynaptic inhibition between CINs are not located on local striatal neurons, but rather on afferent inputs, thereby identifying a long-range GABAergic projection as the principal mediator of CIN feedback inhibition during population activation.

This interpretation is well supported by anatomical and circuit-mapping studies demonstrating that CINs receive substantial extrastriatal innervation. Rabies-based tracing approaches have revealed that CINs integrate inputs from multiple brain regions^34,35^. Among these, inhibitory projections from the external globus pallidus (GPe) emerge as a prominent source of GABAergic input to CINs. In particular, Klug et al. demonstrated that GPe projections directly target CINs and are sufficient to suppress their tonic firing, providing a clear mechanism through which long-range inhibition can shape cholinergic activity.

In parallel, GABAergic projections from the ventral tegmental area (VTA) have been shown to selectively innervate CINs and induce a robust pause in their firing through GABA_A_ receptor-mediated transmission^32^. These findings are particularly notable because they establish that long-range inhibitory inputs can directly reproduce the characteristic pause response of CINs observed in vivo. Together, these studies provide strong evidence that extrastriatal GABAergic pathways are not only anatomically present but functionally capable of exerting powerful inhibitory control over CIN activity.

While our data do not yet identify the precise anatomical origin of the afferent(s) recruited by synchronized CIN activation, they demonstrate that such extrastriatal inhibitory pathways can be engaged via nicotinic signaling, generating a polysynaptic inhibitory loop between CINs. This extends previous work by showing that CINs are not only targets of long-range inhibition, but can actively recruit these inputs through acetylcholine release.

Mechanistically, our interpretation is that β2-containing nAChRs are expressed on the striatal terminals of long-range GABAergic afferents, where they regulate neurotransmitter release. Under this model, synchronized CIN activation leads to widespread acetylcholine release, which activates presynaptic β2-containing nAChRs on GABAergic terminals and promotes GABA release onto CINs. This mechanism closely parallels the well-established role of presynaptic nAChRs in controlling dopamine release from nigrostriatal axons^44,45^, but extends this principle to long-range inhibitory projections.

In summary, our findings reveal a previously unrecognized circuit motif in which CIN population activity recruits extrastriatal GABAergic inhibition through presynaptic nicotinic signaling. This mechanism expands the functional architecture of the striatum beyond local microcircuits, demonstrating that cholinergic interneurons can engage distributed inhibitory networks to regulate their own activity. Such an organization provides a powerful and flexible framework for coordinating striatal dynamics during behavior, linking local cholinergic signaling with long-range circuit control.

### Dopamine exerts a powerful D2-dependent gating of feedback inhibition

Dopamine has long been implicated in regulating the activity of CINs and shaping their characteristic pause responses^10,15,18,19,23^. In particular, D2 receptor signaling modulates CIN excitability and acetylcholine release, and has been proposed to regulate inhibitory interactions within the CIN network^24,25,33,65^

Our results confirm and extend this framework. Activation of D2 receptors with quinpirole nearly abolished feedback inhibition, whereas D1 receptor activation had no detectable effect. Combined D1/D2 agonism produced the same suppression, indicating that this effect is mediated specifically by D2 receptors. This is consistent with Dorst et al. (2020), who showed that D2 receptor activation strongly suppresses polysynaptic CIN–CIN inhibition, an effect attributed to reduced acetylcholine release rather than changes at the GABAergic synapse itself. In contrast, the reduction of feedback inhibition observed following D2 receptor blockade is less straightforward to interpret and likely reflects more complex network-level effects. Together, these findings demonstrate that dopaminergic signaling exerts a strong control over this circuit through D2 receptor activation.

Importantly, despite this strong modulation, the core circuit does not depend on dopamine. Feedback inhibition was fully preserved following 6-OHDA lesions, indicating that β2-nAChR-dependent recruitment of GABAergic inputs operates independently of dopaminergic tone. Dopamine therefore acts as a state-dependent regulator of circuit expression, rather than an essential component of the underlying mechanism.

Functionally, these findings suggest that dopaminergic signaling gates the ability of synchronized CIN activity to recruit inhibition, likely by controlling acetylcholine release and, consequently, the activation of β2-containing nAChRs on downstream elements. This provides a mechanism through which dopamine can dynamically tune cholinergic network inhibition according to behavioral state, while leaving the fundamental circuit architecture intact.

### Functional implications for CIN pause generation and striatal computation

While multiple mechanisms have been proposed to account for the characteristic pause response in CINs, including intrinsic membrane properties, thalamostriatal excitation, and dopaminergic modulation, a unifying circuit mechanism has remained elusive. Our findings provide a framework in which synchronized activation of CINs is intrinsically coupled to delayed inhibition through recruitment of a long-range GABAergic input. In this model, excitatory drive (e.g., thalamostriatal afferents) engages a population of CINs, leading to widespread acetylcholine release. This cholinergic signal, in turn, activates β2-containing nAChRs on GABAergic afferent terminals, triggering GABA release back onto CINs and producing a coordinated suppression of firing. Such a mechanism naturally links excitation and inhibition in sequence, offering a circuit-level substrate for the generation and synchronization of pause responses.

This architecture provides several computational advantages. First, population-dependent recruitment introduces a nonlinear threshold for inhibition, enhancing sensitivity to salient or behaviorally relevant inputs. Second, the involvement of presynaptic nAChRs allows for rapid and temporally precise control of GABA release, consistent with the fast timescale of CIN pause dynamics. Finally, the involvement of long-range inputs allows integration of distributed circuit signals into local striatal processing^1^.

More broadly, these findings highlight a previously underappreciated role for long-range GABAergic projections in shaping striatal microcircuit dynamics, expanding classical models.

## Conclusion

We demonstrate that synchronized activation of striatal cholinergic interneurons recruits a powerful, β2-nAChR-dependent inhibitory pathway originating from extrastriatal GABAergic afferents. This circuit provides a mechanistic link between population activity and coordinated inhibition, is strongly gated by dopamine, and operates independently of local striatal interneurons. These findings redefine the circuit basis of cholinergic signaling in the striatum and reveal a central role for long-range inhibitory inputs in controlling striatal network dynamics.

## Acknowledgements

We thank Fulva Shah for excellent technical assistance and expertise in the management and maintenance of transgenic mouse colonies. This work was supported by a Michael J. Fox Foundation for Parkinson’s Research Grant (MJFF-022900; to M.A.), a NARSAD Young Investigator Award from the Brain & Behavior Research Foundation (Grant ID: 27897; to M.A.), a Busch Biomedical Research Grant (to M.A.), and a Royal Society Research Grant (RG\R2\232186; to M.A.). Additional support was provided by the National Institute of Neurological Disorders and Stroke (NINDS) grant R01 NS034865 (to J.M.T.).

## Author contributions

M.A. designed the study, obtained funding, collected and analyzed data, and wrote the manuscript with input from all authors. S.K. and E.B.G. collected and analyzed data and participated in experimental design. J.M.T. obtained funding and participated in study design.

The authors declare no competing interests.

## Materials & Correspondence

Correspondence and requests for materials should be addressed to Maxime Assous.

## Methods

### Resource availability

Further information and requests for resources and reagents should be directed to and will be fulfilled by the lead contact, Maxime Assous. This study did not generate new unique reagents. All data supporting the findings of this study are available from the corresponding author upon reasonable request. Custom MATLAB scripts used for analysis are available from the lead contact upon request. Additional information required to reanalyse the data reported in this study is available upon request.

### Animals

All procedures were performed in accordance with the UK Animals (Scientific Procedures) Act 1986 and approved by institutional ethical review committees. Experimental procedures were also consistent with the National Institutes of Health Guide for the Care and Use of Laboratory Animals.

Adult male and female mice aged 3–8 months were used for all experiments. The following transgenic mouse lines were employed: ChAT-ChR2-EYFP (Tg(Chat-COP4*H134R/EYFP,Slc18a3)6Gfng/J; The Jackson Laboratory, Stock No: 014546), TH-Cre (Tg(TH-Cre)12Gsat; GENSAT), Htr3a-Cre (Tg(Htr3a-Cre)NO152Gsat; MMRRC), SST-Cre (Sst-IRES-Cre; The Jackson Laboratory, Stock No: 013044), VGAT-Cre (Slc32a1tm2(cre)Lowl/J; The Jackson Laboratory, Stock No: 016962), VGlut2-Cre (Slc17a6tm2(cre)Lowl/J; The Jackson Laboratory, Stock No: 016963), and β2-floxed mice (Chrnb2^fl/fl; Chrnb2tm1.1Mccl; MGI:5702916; provided by Dr. Crair, Yale University). Double transgenic animals were generated by crossing ChAT-ChR2-EYFP mice with the appropriate Cre-driver lines, and only mice positive for both transgenes were used for experiments. For conditional knockout experiments, ChAT-ChR2-EYFP::β2^fl/fl mice were generated by crossing ChAT-ChR2-EYFP mice with β2^fl/fl mice. Mice were housed in groups of up to four per cage under a 12 h light/dark cycle with ad libitum access to food and water.

### Stereotaxic viral injections

Mice were anesthetized with isoflurane (1–2.5% in O₂, 1 L/min) and placed in a stereotaxic frame (Kopf). Bupivacaine was applied subcutaneously at the incision site for local analgesia. Viral vectors were delivered through glass micropipettes using a Nanoject II Auto-Nanoliter Injector (Drummond Scientific, Cat. No. 3-000-204).

The dorsal striatum was targeted using coordinates relative to bregma of AP +0.6 mm, ML ±1.9 mm, and DV −2.5 mm and −3.0 mm from the brain surface. For conditional deletion of β2-nAChRs in striatal neurons, ChAT-ChR2-EYFP::β2^fl/fl mice received unilateral injections of AAV2-hSyn-mCherry-Cre (University of North Carolina Vector Core, Cat. No. AV6145B) and contralateral injections of AAV2-hSyn-mCherry (UNC Vector Core, Cat. No. AV5033E) as control. Viruses were delivered at a total volume of 1.66 μL per injection site at a rate of 13.8 nL every 5 s. Recordings were performed in both hemispheres of the same animal.

For intersectional experiments designed to record from CINs lacking ChR2 expression, ChAT-Cre mice were injected bilaterally with AAV5-hSyn-Con/Foff-ChR2(H134R)-eYFP (UNC Vector Core, Cat. No. AV8475) together with a mixture of AAV5-EF1a-DIO-FLPo-WPRE-hGHpA (Addgene, Cat. No. 87306) and AAV5-EF1a-fDIO(FRT)-mCherry (Addgene, Cat. No. 114471). This Cre-ON/Flp-OFF strategy enables expression of ChR2 in CINs that undergo Cre recombination but not Flp recombination, while CINs expressing both recombinases undergo Flp-mediated recombination that inactivates ChR2 and induces mCherry expression. A total volume of 1.5 μL of the Con/Foff virus was injected per hemisphere at a rate of 13 nL every 4 s, followed by 200 nL of the Flp-dependent viral cocktail (mixed 1:1) delivered at 2.3 nL every 5 s. Recordings were restricted to mCherry⁺/ChR2-eYFP⁻ CINs.

For retrograde deletion of β2-nAChRs from striatal afferents, ChAT-ChR2-EYFP::β2^fl/fl mice received bilateral injections of AAV-retro-pgk-Cre (Addgene, Cat. No. 24593) together with AAV-Flex-tdTomato (Addgene, Cat. No. 28306) into the dorsal striatum. In the contralateral hemisphere, control injections consisted of AAV-retro-EF1a-FLPo (Addgene, Cat. No. 55637) combined with AAV5-EF1a-fDIO(FRT)-mCherry (Addgene, Cat. No. 114471). Retrograde transport of Cre recombinase induced excision of the floxed β2 subunit selectively in neurons projecting to the striatum, while recorded CINs themselves did not undergo recombination. A total volume of 1.5 μL per hemisphere was delivered at a rate of 9.3 nL every 4s. Fluorescent reporter viruses were diluted 1:100 before injection to minimize overexpression.

For optogenetic inhibition experiments, AAV5-EF1α-DIO-eNpHR3.0-mCherry was injected into the striatum of VGAT-Cre, SST-Cre, Htr3a-Cre, or TH-Cre mice to enable cell-type-specific expression of halorhodopsin (UNC Vecor Core, AV4832E). For activation of thalamostriatal afferents, VGlut2-Cre mice were injected with AAV5-EF1α-DIO-ChR2-eYFP (UNC Vector core, AV4378P) into the parafascicular nucleus using coordinates AP −2.3 mm, ML ±0.75 mm, DV −3.3 mm, with a total volume of 400 nL.

Following viral delivery, the pipette was left in place for 10 min before slow retraction. Animals received postoperative analgesia (ketoprofen and buprenorphine) and were allowed to recover for at least 4 weeks prior to experiments.

### 6-OHDA lesions

To induce dopaminergic lesions, ChAT-ChR2-EYFP mice were injected unilaterally in the substantia nigra pars compacta with 6-hydroxydopamine (6-OHDA; 3.5 mg/mL in 0.1% ascorbic acid; Sigma-Aldrich, Cat. No. H4381), while the contralateral hemisphere received saline as control. Injections were performed at coordinates AP −2.7 mm, ML ±1.35 mm, DV −4.2 mm relative to bregma. A total volume of 1.3 μL was delivered at 13.8 nL every 5 s.

### Slice preparation

Mice were deeply anesthetized with ketamine (80 mg/kg) and xylazine (20 mg/kg) and transcardially perfused with ice-cold oxygenated NMDG-based solution containing (in mM): 103 NMDG, 2.5 KCl, 1.2 NaH₂PO₄, 30 NaHCO₃, 20 HEPES, 25 glucose, 101 HCl, 10 MgSO₄, 2 thiourea, 3 sodium pyruvate, 12 N-acetyl cysteine, and 0.5 CaCl₂, equilibrated with 95% O₂ and 5% CO₂ (pH 7.2–7.4). Brains were rapidly removed and sectioned into 300 μm parasagittal slices using a vibratome (Leica VT1200S).

Slices were recovered in oxygenated NMDG solution at 35 °C for 5 min and then transferred to artificial cerebrospinal fluid (ACSF) containing (in mM): 124 NaCl, 2.5 KCl, 1.2 NaH₂PO₄, 26 NaHCO₃, 1 MgCl₂, 2 CaCl₂, 10 glucose, and 3 sodium pyruvate (pH 7.2–7.4, continuously bubbled with 95% O₂ and 5% CO₂) and maintained at 25 °C until recording. During recordings, slices were perfused with ACSF at 32–34 °C at a rate of 2–4 mL/min.

### Electrophysiology

Whole-cell recordings were performed using borosilicate glass pipettes pulled using a vertical puller (Narishige PP-83). Pipette resistance ranged from 3 to 5 MΩ. Voltage-clamp recordings were performed at a holding potential of −45 mV unless otherwise specified.

The CsCl-based internal solution used for voltage-clamp recordings contained (in mM): 125 CsCl, 0.1 EGTA, 10 HEPES, 2 MgCl₂, 4 Na₂ATP, and 0.4 Na₂GTP, supplemented with 5 mM QX-314 (Tocris) and 0.2% biocytin (Sigma-Aldrich). The K-gluconate-based internal solution used for current-clamp recordings contained (in mM): 130 K-gluconate, 10 KCl, 2 MgCl₂, 10 HEPES, 4 Na₂ATP, and 0.4 Na₂GTP.

Signals were recorded using a Multiclamp 700B amplifier (Molecular Devices), digitized at 20 kHz using a CED Micro1401 Mk II (Cambridge Electronic Design), and acquired using Signal software (version 5; Cambridge Electronic Design).

### Optogenetic stimulation

Optogenetic stimulation was delivered using a high-power multi-colour LED (LZC-R0H100-0000, Mouser Electronics), coupled to the microscope optical path (specific optical configuration not specified; please provide if available). Channelrhodopsin-2 was activated using 2 ms blue light pulses (450 nm). Halorhodopsin (eNpHR3.0) was activated using 590 nm light pulses of 700 ms duration, beginning 200 ms prior to ChR2 stimulation. Stimuli were delivered every 30 s.

### Pharmacology

Pharmacological agents were bath-applied in ACSF. Bicuculline (10 μM; Sigma-Aldrich) was used to block GABA_A receptors. Dihydro-β-erythroidine hydrobromide (DHβE; 1 μM; Tocris) was used to block β2-containing nicotinic acetylcholine receptors. CNQX (10 μM; Tocris) and D-APV (10 μM; Tocris) were used to block AMPA and NMDA receptors, respectively. Tetrodotoxin (TTX; 1 μM; Sigma-Aldrich) was used to block voltage-gated sodium channels and isolate the photocurrent.

Dopaminergic modulation was examined using SCH-23390 (10 μM; Sigma-Aldrich) and SKF-38393 (10 μM; Sigma-Aldrich) for D1 receptor blockade and activation, respectively, and sulpiride (5 μM; Tocris) and quinpirole (10 μM; Tocris) for D2 receptor blockade and activation, respectively. Drugs were applied following a stable baseline period of 20 trials (10 minutes).

### Histology and imaging

Brains were fixed by transcardial perfusion with 4% paraformaldehyde and post-fixed overnight. Coronal sections (50 μm) were obtained using a vibratome (Vibratome 1500 Sectioning System). For tyrosine hydroxylase (TH) immunohistochemistry, sections were incubated overnight at 4 °C with rabbit anti-TH primary antibody (Millipore, 1:1500) in a solution containing 10% normal donkey serum, 0.5% Triton X-100, and 2% bovine serum albumin in PBS. Following washes, sections were incubated for 5 h at room temperature with Alexa Fluor 594-conjugated secondary antibody (1:250, Invitrogen).

Biocytin-filled neurons were visualized using streptavidin-Alexa Fluor 405 (Thermo Fisher Scientific, 1:1000) and anti-GFP antibody (Invitrogen, 1:1000). Sections were mounted using Vectashield mounting medium (Vector Laboratories).

Images were acquired using an Olympus Fluoview FV1000 confocal microscope equipped with 10× and 40× objectives. Images were processed using ImageJ.

### Quantification and statistical analysis

Electrophysiological data were analyzed using Signal (version 5; Cambridge Electronic Design) and Axograph (version 1.3.4). To isolate synaptic currents from overlapping ChR2-mediated photocurrents, a custom MATLAB script was used to fit and subtract a biexponential function of the form:

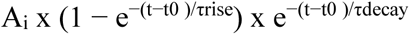

where *τ rise* and *τ decay* represent the rise and decay time constants, respectively, and Ai represents amplitude scaling. This approach enabled extraction of synaptic latency, peak amplitude, and charge transfer independent of the photocurrent.

Statistical analyses were performed using GraphPad Prism 9 (GraphPad Software). Data were tested for normality using the Shapiro–Wilk test. Two-group comparisons were performed using paired or unpaired two-tailed t-tests as appropriate, while multiple comparisons were analyzed using one-way or two-way ANOVA followed by Tukey’s post hoc test. Statistical significance was defined as p < 0.05.

Data are presented as mean ± SEM unless otherwise stated. The variable n represents the number of cells, and N represents the number of animals. Analyses were performed blind to experimental condition where feasible.

---

**Figure S1.**
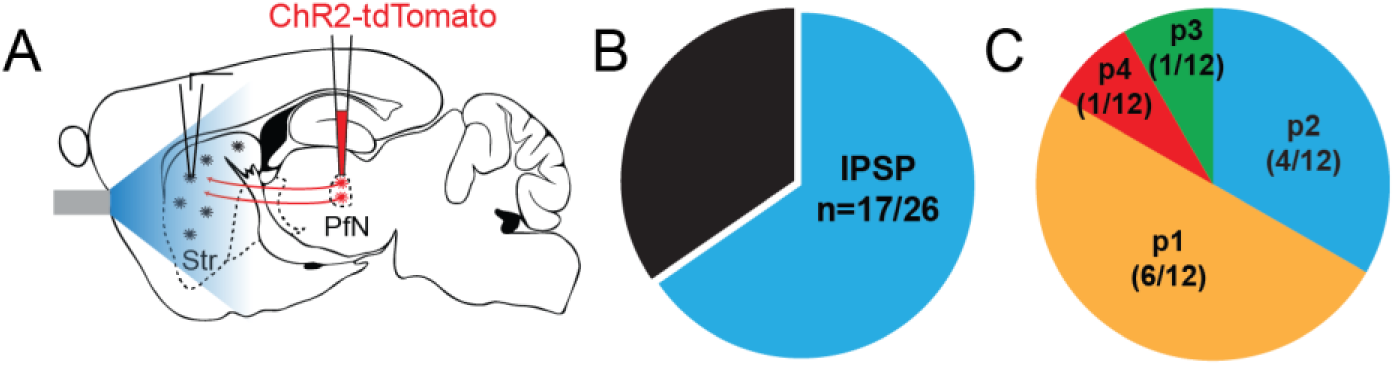
(Related to Figure 1). Proportion of CINs exhibiting thalamostriatal-evoked feedback inhibition. (A) Schematic illustrating the experimental strategy used to activate thalamostriatal afferents. (B) Proportion of recorded CINs exhibiting optogenetically-evoked inhibitory postsynaptic potentials/currents (IPSPs/IPSCs) following PfN stimulation (17/26 cells). (C) Distribution of inhibitory responses according to the pulse number within a train of five optical stimulations delivered at 20 Hz. p1–p5 indicate IPSCs/IPSPs first appearing after the first, second, third, fourth, or fifth stimulation pulse, respectively.

**Figure S2.**
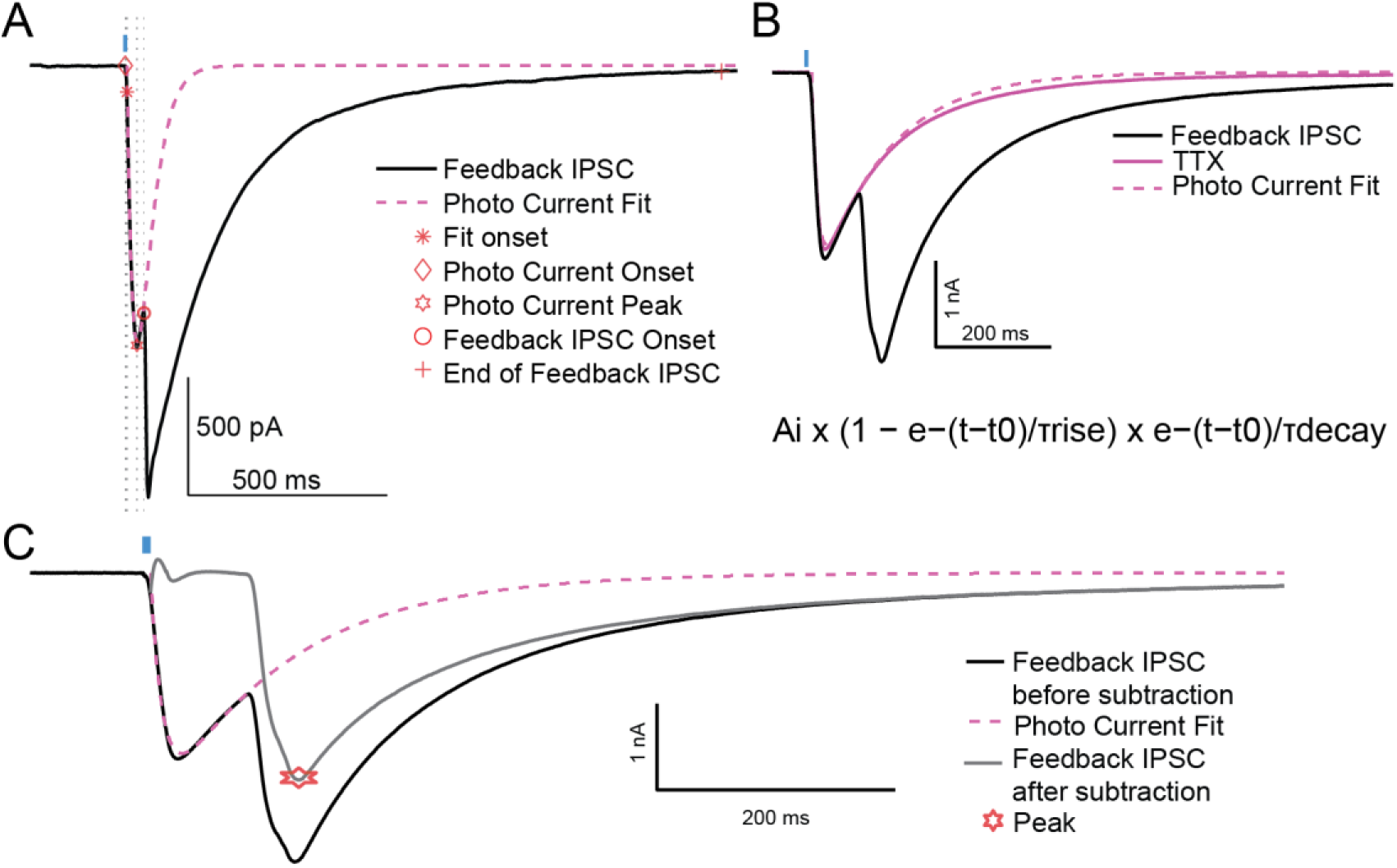
(Related to Figure 2). Validation of the biexponential fitting approach used to isolate feedback IPSCs from overlapping ChR2-mediated photocurrents in cholinergic interneurons. (A) Representative voltage-clamp recording from a ChR2-expressing cholinergic interneuron (CIN) showing the overlap between the direct optogenetically-evoked photocurrent and the delayed feedback IPSC. A custom MATLAB-based biexponential fitting procedure was used to model the photocurrent component according to the function: Ai × (1−e−(t−t0)/τrise) × e−(t−t0)/τdecay. The fit identified the onset and peak of the photocurrent as well as the onset and duration of the feedback IPSC. (B) Validation of the fitting procedure following bath application of tetrodotoxin (TTX, 1 μM), which abolished synaptic transmission and isolated the photocurrent component. The biexponential model accurately reproduced the remaining photocurrent waveform. (C) Representative trace illustrating isolation of the feedback IPSC following subtraction of the fitted photocurrent from the raw recording. This analytical approach was subsequently applied to all recordings to quantify feedback IPSC onset latency, peak amplitude, decay kinetics, and charge transfer.

**Figure S3.**
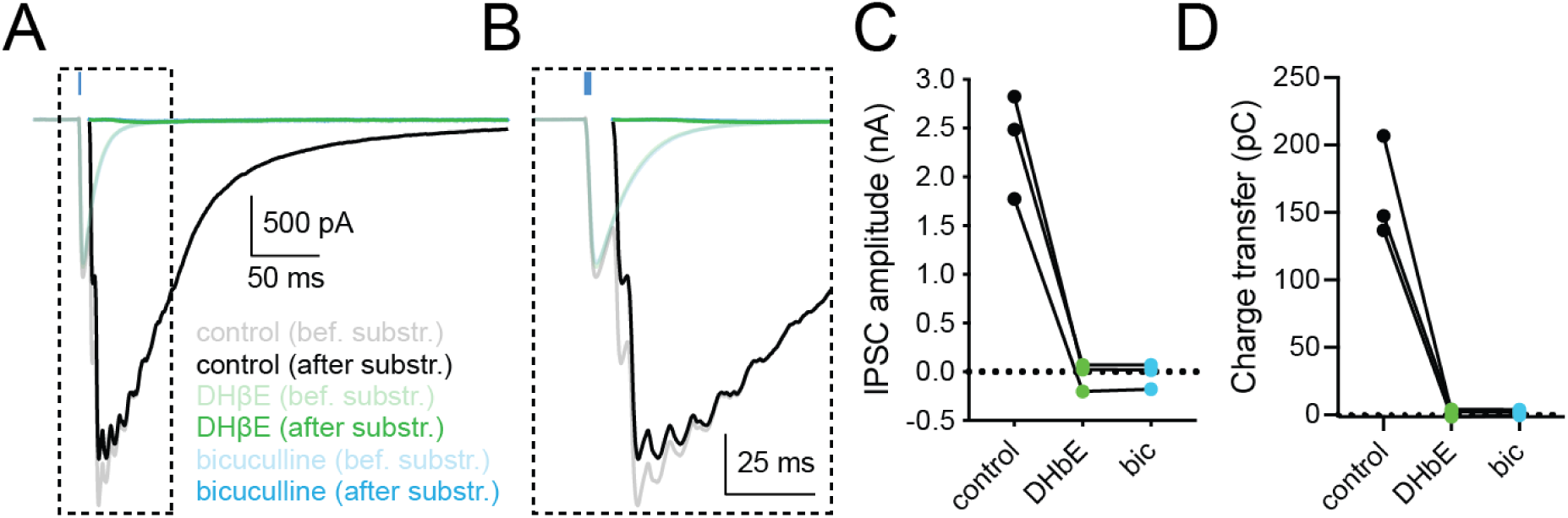
(Related to Figure 2). GABAA receptor blockade does not further reduce feedback inhibition following β2-nAChR antagonism. (A) Representative voltage-clamp recordings from ChR2-expressing cholinergic interneurons (CINs) showing optogenetically-evoked feedback IPSCs (before and after photocurrent subtraction) under control conditions, following application of the β2-containing nAChR antagonist DHβE (1 μM, green), and after subsequent addition of bicuculline (10 μM, blue). (B) Expanded traces illustrating the isolated feedback IPSCs after subtraction of the fitted photocurrent under each condition. Application of DHβE nearly abolished the feedback IPSC, and subsequent GABAA receptor blockade with bicuculline produced no additional reduction. (C) Quantification of feedback IPSC peak amplitude under control conditions, after DHβE, and following bicuculline application in the continued presence of DHβE. (D) Quantification of IPSC charge transfer across conditions. Each dot represents one recorded cell.

**Figure S4.**
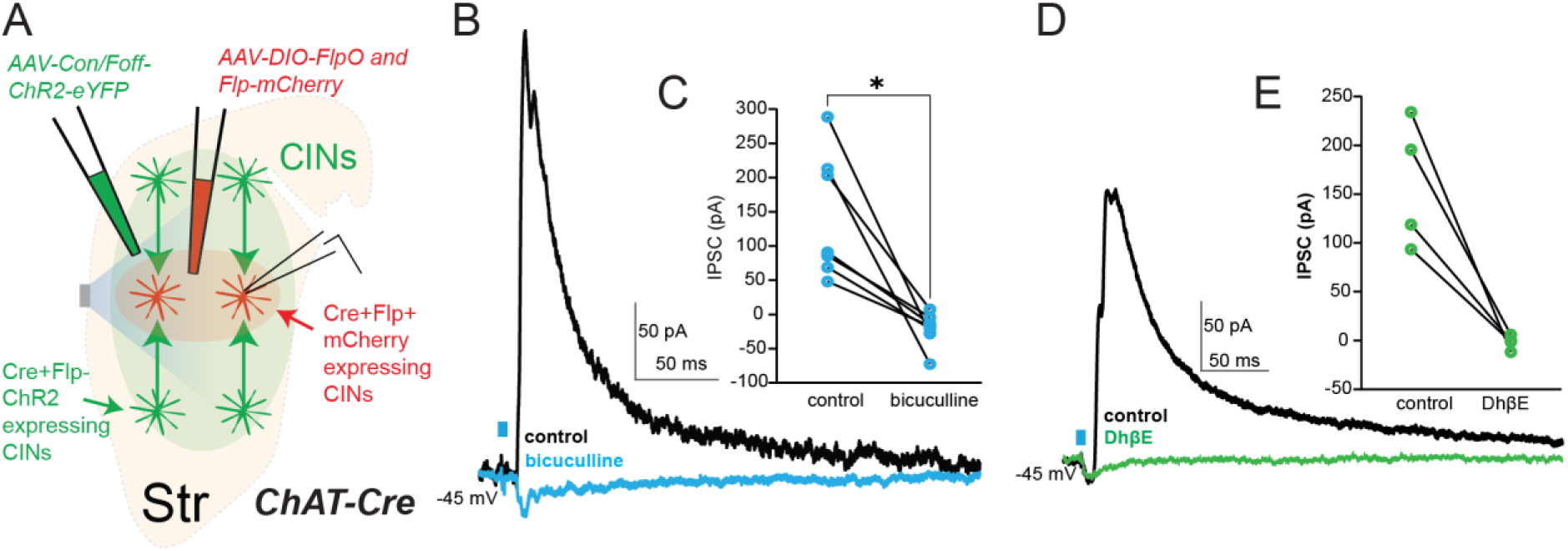
(Related to Figure 3). Intersectional strategy to record from ChR2-negative cholinergic interneurons during optogenetic activation of neighboring CINs. (A) Schematic of the intersectional viral strategy used to generate two spatially intermingled but genetically distinct CIN populations in ChAT-Cre mice. A Cre-dependent Con/Foff-ChR2-eYFP construct was co-injected with AAV-DIO-FlpO and Flp-mCherry into the dorsal striatum. CINs expressing Cre alone retained ChR2-eYFP expression and were optogenetically stimulated, whereas CINs co-expressing Cre and virally delivered Flp underwent Flp-mediated inversion of the ChR2 cassette, resulting in mCherry-positive / ChR2-negative CINs used for whole-cell recordings. (B) Representative voltage-clamp recordings obtained from ChR2-negative/mCherry-positive CINs held at −45 mV during optical stimulation of neighboring ChR2-positive CINs. Optogenetic stimulation evoked robust IPSCs in recorded CINs under control conditions, which were abolished by the β2-containing nAChR antagonist DHβE (1 μM). (C) Quantification of IPSC amplitude before and after DHβE application. (D) Representative voltage-clamp recordings from ChR2-negative CINs showing that optogenetically-evoked IPSCs were abolished by the GABAA receptor antagonist bicuculline (10 μM). (E) Quantification of IPSC amplitude under control conditions and following bicuculline application. These experiments confirm that inhibitory coupling between CINs can be observed independently of direct ChR2-mediated photocurrents and depends on β2-containing nAChRs and GABAA receptor-mediated transmission. Each dot represents one recorded cell. P < 0.05.

**Figure S5.**
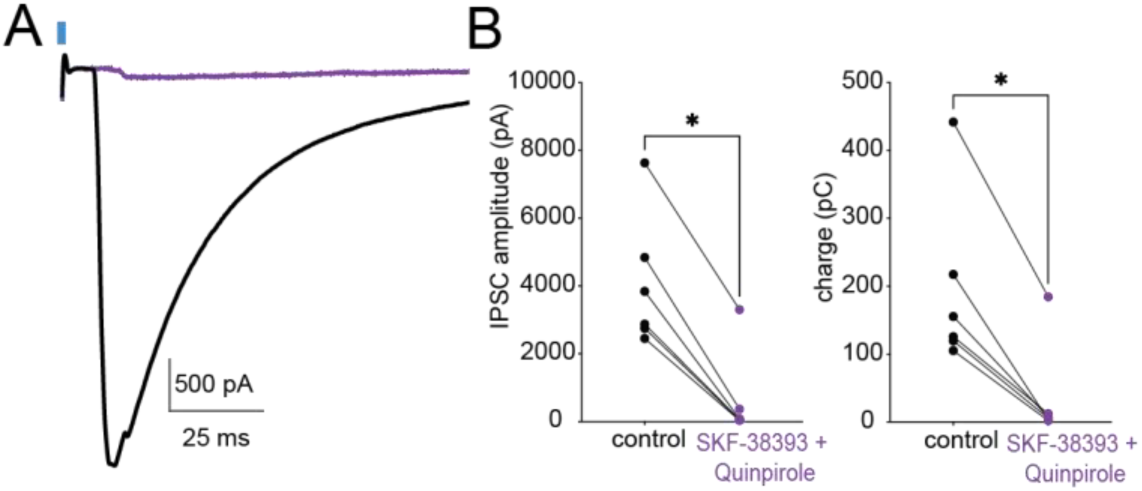
(Related to Figure 5). Combined D1 and D2 receptor activation suppresses feedback inhibition between cholinergic interneurons. (A) Representative voltage-clamp recordings from ChR2-expressing CINs showing optogenetically-evoked feedback IPSCs under control conditions and following co-application of the D1 receptor agonist SKF-38393 (10 μM) and the D2 receptor agonist quinpirole (10 μM). Combined dopamine receptor activation strongly reduced the amplitude of feedback inhibitory responses. (B) Quantification of IPSC peak amplitude and charge transfer before and after co-application of SKF-38393 and quinpirole. Each line represents one recorded cell. *P < 0.05.

**Figure S6.**
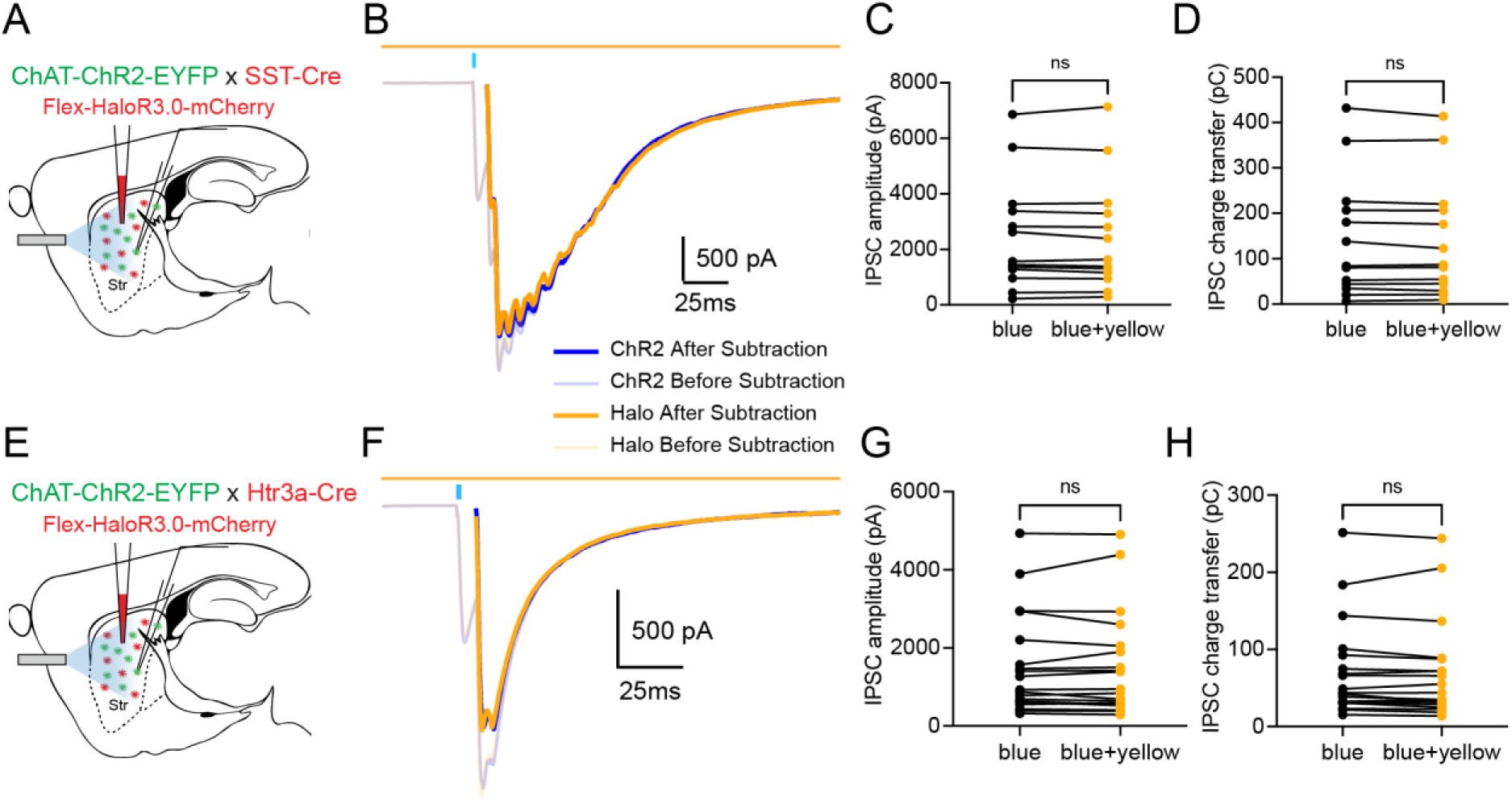
(Related to Figure 6). Optogenetic inhibition of Htr3a- and SST-expressing interneurons does not alter feedback inhibition between cholinergic interneurons. (A) Schematic of the experimental strategy used to inhibit somatostatin-expressing interneurons. ChAT-ChR2-eYFP × SST-Cre mice received striatal injections of a Cre-dependent HaloR3.0-mCherry AAV, enabling optogenetic silencing of SST interneurons during activation of CINs. (B) Representative voltage-clamp recordings from CINs showing optogenetically-evoked feedback IPSCs during blue light stimulation alone and during simultaneous blue + yellow light stimulation to inhibit SST interneurons. Traces are shown before and after photocurrent subtraction. (C,D) Quantification of IPSC peak amplitude (C) and charge transfer (D) under both conditions. Optogenetic inhibition of SST interneurons did not significantly affect feedback inhibition between CINs. (E) Schematic of the double optogenetic strategy used to test the contribution of Htr3a-expressing interneurons to feedback inhibition between CINs. ChAT-ChR2-eYFP × Htr3a-Cre mice received striatal injections of a Cre-dependent HaloR3.0-mCherry AAV, enabling optogenetic inhibition of Htr3a interneurons during blue light activation of ChR2-expressing CINs. (F) Representative voltage-clamp recordings from CINs showing optogenetically-evoked feedback IPSCs during blue light stimulation alone and during simultaneous blue + yellow light stimulation to inhibit Htr3a-expressing interneurons. Traces are shown before and after photocurrent subtraction. (G,H) Quantification of IPSC peak amplitude (G) and charge transfer (H) under blue light stimulation alone or during simultaneous inhibition of Htr3a interneurons. Optogenetic suppression of Htr3a interneurons did not significantly alter feedback inhibition between CINs.

